# The medial entorhinal spatial map is built on excitatory-inhibitory network motifs defined by their functional cell type and theta modulation

**DOI:** 10.64898/2026.07.02.735850

**Authors:** Pauline Kerekes, Marius Bauza, Julija Krupic

**Affiliations:** UK Dementia Research Institute, University College London, London, UK; Sainsbury Wellcome Centre for Neural Circuits and Behaviour, University College London, London, UK; Department of Physiology, Development and Neuroscience, University of Cambridge, Cambridge, UK

## Abstract

The medial entorhinal cortex (mEC) contains functionally specific neurons crucial for spatial memory and navigation, including grid, border, spatial, head direction, and cue cells. However, how these neurons interact to build spatial maps remains unclear. Here, using simultaneous recordings from hundreds of functionally defined mEC neurons in mice navigating virtual tracks with varying numbers of visual landmarks, we uncovered connectivity motifs that define the mEC functional network architecture and link it to the principles governing allocentric map formation. We found that connectivity between cells was cell-type-specific and grouped into subnetworks based on their function and theta modulation, with theta-modulated connections dominating over non-theta. Functionally distinct neurons preferentially connected to their own type, and their interactions were coordinated by a shared inhibitory pool of interneurons, with a higher proportion of excitatory-to-inhibitory connections than between excitatory cells. This was accompanied by the number of fields formed across all mEC cell types increasing sublinearly with the number of available cues. Grid cells showed the strongest relative connectivity to interneurons regardless of their theta modulation, linking otherwise largely isolated theta and non-theta streams. Grid cells were also least likely to form a field in response to cues. Together, these findings reveal an mEC network architecture organised by cell type and theta dependency, in which inhibitory interactions, with grid cells as the strongest hub, play a central role in building the mEC allocentric map.

## Main

Functionally specialized neurons, such as grid^1^, head direction^2^, border^3^, cue^4^ and spatially tuned cells^5^, constitute the core elements of the medial entorhinal (mEC) spatial map^6,7^. Collectively, these cells integrate diverse information about landmarks, distances, and the direction of travel to construct a cohesive spatial map. However, the principles governing the field formation of functionally distinct mEC cells in response to visual cues have not been characterized. Because mEC excitatory cells are connected via a rich inhibitory cell network^8,9^, interneurons may play a key role in implementing coordination, similar to recent findings in the hippocampus proper^10,11^. However, unlike hippocampal place cells, mEC principal cells have markedly distinct functional properties and therefore may follow different rules of organization. Indeed, different inhibitory molecular cell types in the mEC have different effects on functionally distinct principal cells^8,12,13^. Finally, given the role of theta oscillations in memory and navigation^6,14^, they may be expected to play an important role in network coordination by providing temporal windows for local circuit interactions^14–16^.

Here, we used high-throughput Neuropixels 2.0 recordings^17^ in mice navigating virtual reality (VR) environments populated with different numbers of cues to simultaneously record from hundreds of functionally defined mEC cells to examine how their neural network architecture may underlie the construction of mEC spatial maps. We revealed specific neural network motifs that structure mEC cell interactions and may support cue-dependent field formation.

## Results

To study how medial entorhinal (mEC) cells collectively build a visual scene, we used Neuropixels 2.0 to record mEC grid cells^1^ (GC, n=944 across three grid modules), head-direction cells^2^ (HD, n=155), border cells^3^ (BC, n=186), spatially tuned cells^5^ (SC, n=760) and cue cells^4^ (CC, n=1115) in mice navigating in virtual reality (VR) environments (Fig. 1a-c, and Extended Data Figs. 1-4). The anatomical mapping revealed that different functional cell types were highly interspersed (Extended Data Fig. 5).

**Fig. 1:**
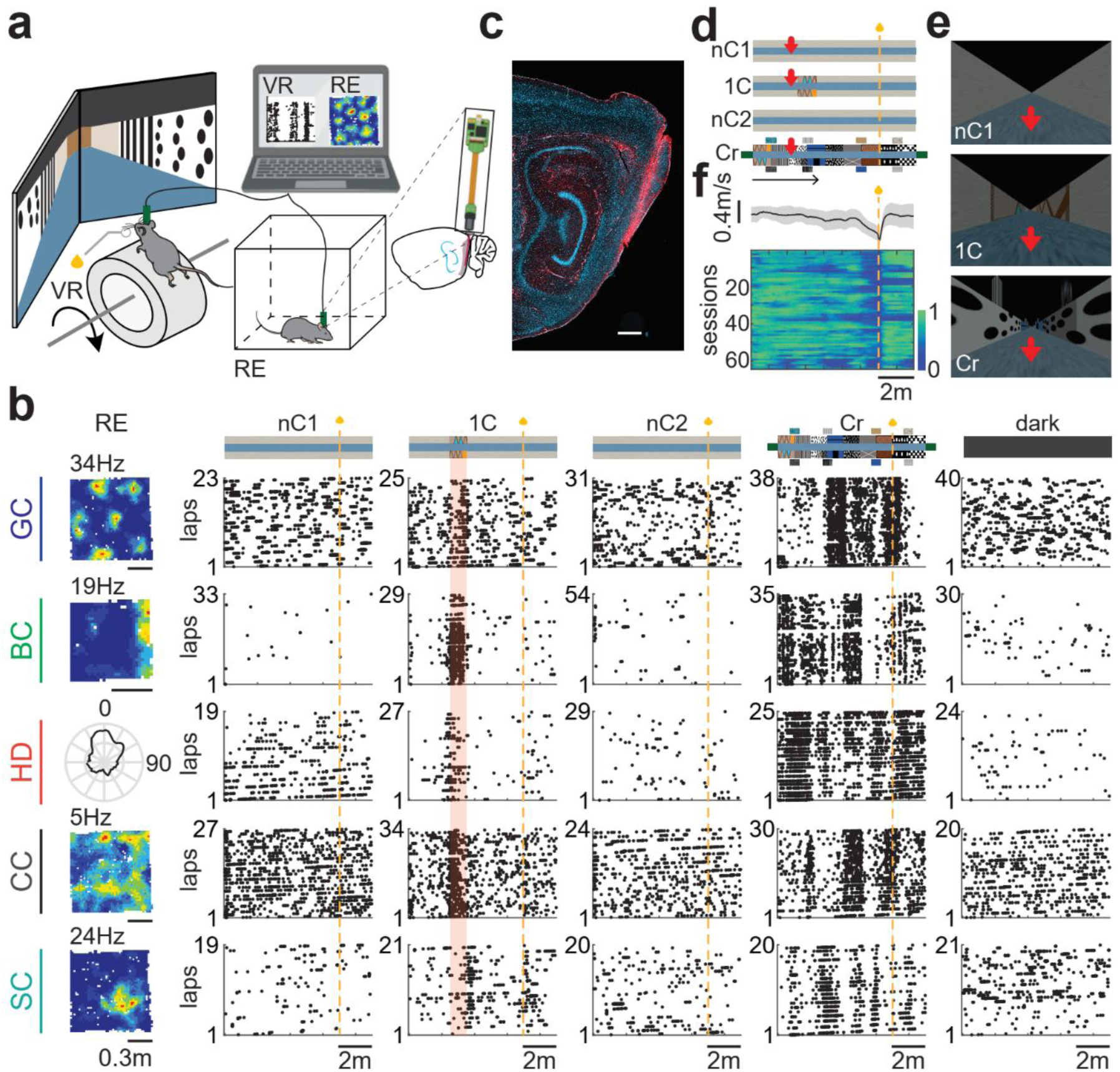
Mice navigating in virtual reality environments coupled with high-throughput neural recordings in the medial entorhinal cortex. **a**, Schematics of a recording session. Left, mice first ran in the virtual reality (VR) tracks followed by a trial in a real enclosure (RE). Right, Neuropixels 2.0. implanted in the left mEC. **b**, Firing rate maps (top left: peak firing rate) in the RE and raster plots in the first and second trials of no cue (nC1 and nC2), one-cue (1C), cue rich (Cr) and dark **(d)** virtual tracks of six example cells belonging to different cell types (GC, grid cell; BC, border cell, HD, head-direction cell; CC, cue cell; SC, entorhinal spatially tuned cell). In all raster plots, the golden dashed line and the shaded area indicate the reward at 7m and the cue site, respectively. c, Implant location of the Neuropixels probe in the mEC. Red: anti-GFAP staining; blue: DAPI staining. d, Top view of the virtual tracks. The horizontal and red arrows indicate respectively the direction of movement and the position of the screenshots shown in **(e)**. e, Screenshots of the VR as seen by the animal. **f**, Bottom, running speed as a function of the position (5cm per bin) for individual cue-rich (Cr) trials. Each heat map is normalized by its maximum speed. Top, mean±s.d. of the individual trials running speed profiles.

Neural activity was recorded in mice navigating in nine-meter VR linear tracks with no cues (nC), with a single (1C), or multiple cues (Cr, cue-rich), and with a fixed reward at seven meters from the start (Fig. 1d). Mice were teleported back to the start location once they reached the end of the track, with a 2-second dark inter-lap period to signal the end of the lap. The VR session followed an A-B-A-C experimental design, with a no-cue track as a baseline (A), a one-cue track as a probe (B), and a cue-rich track as a final trial (C). The VR session was followed by a final recording in a square real enclosure (RE, 1m x 1m or 0.6m x 0.6m) to characterize the functional cell types recorded in the VR. The trials in VR were separated by a 2-minute dark inter-trial interval, during which the VR screens displayed a dark, homogeneous background, and no reward was given to the animal. The no-cue track consisted of identical homogeneous walls and floor with a random granular visual texture (Fig. 1d-e, Extended Data Video 1) to ensure that the mouse could only use self-motion cues (e.g., efference copy and optic flow) and/or time counting to estimate its location. The second track was identical to the first, with an additional one-meter-long proximal cue displayed on both walls, 2.5 meters from the start (Fig. 1d-e and Extended Data Video 2). There was no wall at the end of the no-cue and one-cue tracks. The final VR environment was a cue-rich track with nine one-meter-wide cues tiling the walls, and four pairs of distal cues positioned at 1, 3, 6, and 8 meters from the start (Fig. 1d-e and Extended Data Video 3). Prior to the A-B-A-C session, mice were trained on the cue-rich track for 4-6 days until they reliably reduced their running speed 1-1.5m before the reward (Fig. 1f). This deceleration was used as a benchmark to assess self-localization in all environments in which the mice were tested.

### No spatial fields formed away from the boundaries without visual cues

First, we investigated how different cell types constructed the spatial map in the absence of visual landmarks in the no-cue VR track. To assess the level of spatial activity, we calculated the number of spatial fields present along the track. The fields were defined as a group of spikes consistently fired on at least 50% of the visits of a specific track location (Extended Data Fig. 6). We found that, irrespective of functional type, most cells did not form stable fields along the track away from the boundaries, with fields forming almost exclusively at the start of the track (Fig. 2a-c, and Extended Data Fig. 7a). The number of cells that formed fields was consistently higher than in complete darkness trials in line with previous reports^18,19^ (Fig. 2a-b; no cue vs. dark track: BC: 32/186 vs. 0/186; CC: 132/1115 vs. 10/1115; HD: 20/155 vs. 0/155; GC: 44/944 vs.10/944; SC: 61/760 vs. 4/760; permutation test: *P*=0). We hypothesized that these fields may have been driven by changes in illumination during the transition between the no-cue environment and the dark inter-lap segment. Indeed, we observed that with the exception of grid cells, all functional cell types consistently showed an increase in firing rate as the animal went in and out of the dark inter-lap segment (Fig. 2d). Grid cells peaked their firing only at the transition between no-cue track and dark interval, suggesting that they either underwent neural adaptation to the visual contrast or they may have signaled the end of the lap akin to the ‘event boundary’ responses described in humans^20^.

**Fig. 2:**
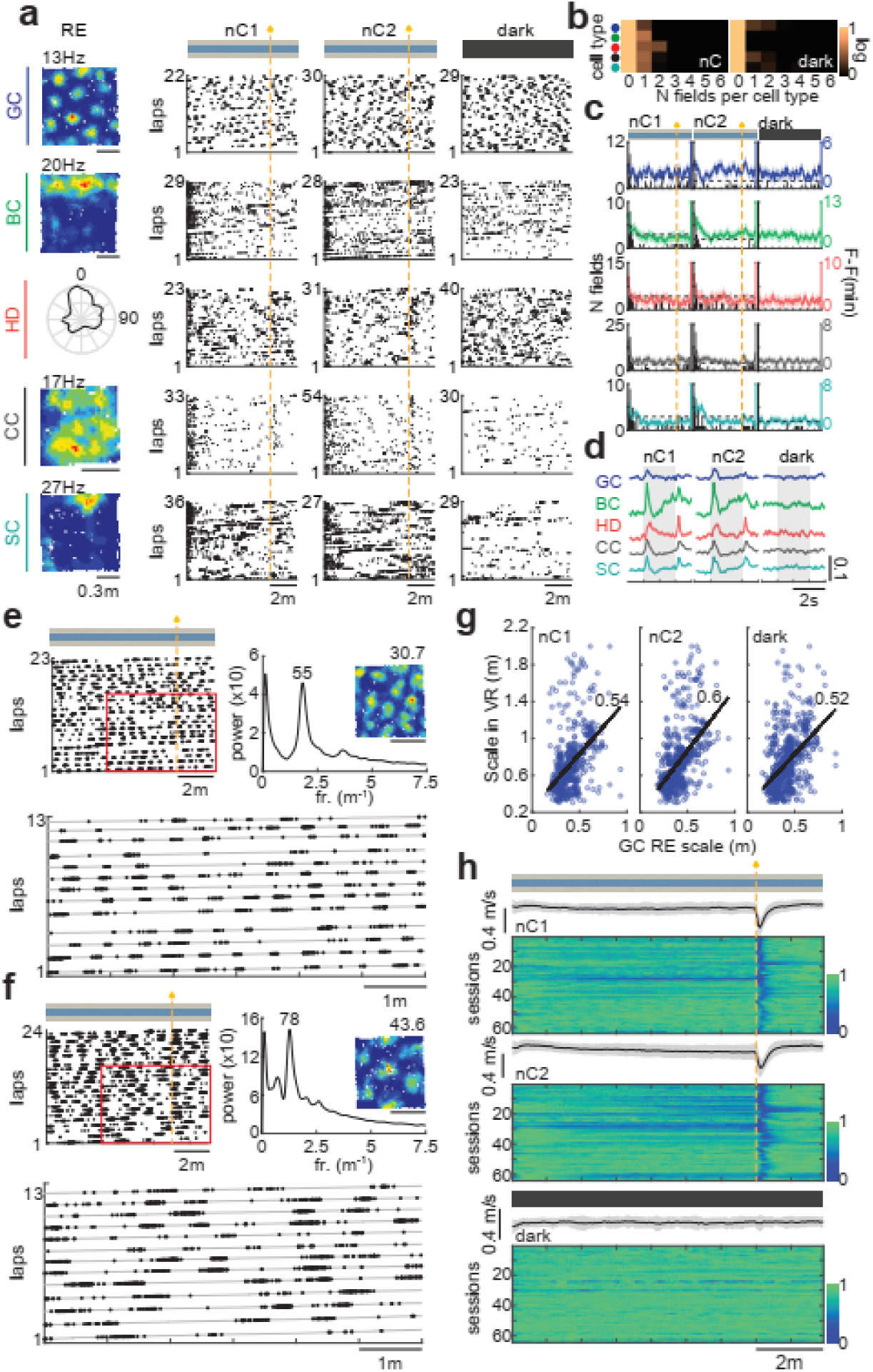
Absence of spatial fields away from the boundaries without visual cues. **a**, Typical rate maps (top left: peak firing rate) and raster plots of different functional cell types in the RE and in the first and second trials of no-cue (nC1 and nC2) and the dark virtual tracks. **b**, Distribution of the number of fields formed per cell for each cell type; color code as shown in (a). All pixel values have been converted to base-10 logarithms for display. **c**, Field count per spatial bin (20cm per bin) overlaid with the mean ± s.e.m. of the smooth firing rate (Hz, aligned by subtracting its minimum value) per spatial bin (5cm per bin) in nC1, nC2, and dark tracks. The gray dashed line indicates the chance level for the field distribution. **d**, The average firing rate (100ms bin) during the dark inter-lap interval (grey shade). Each trace is aligned by subtracting its average value. **e**, A typical grid cell showing periodic activity in the no-cue track. Left: raster plot of the spikes as a function of the VR track position. Bottom: magnified view of the red inset above. Right: the corresponding FFT power spectrogram as a function of the spatial frequency showcasing a prominent peak at a spatial period of 55 cm, which was ∼1.8 times the grid scale (cm) in RE (scale: top right corner of the rate map). Scale bar = 50 cm. **f**, Same as (a) for a grid cell with a larger scale. **g**, Distribution of grid scale in RE and their corresponding scales in VR measured in the first (nC1, left) and second (nC2, middle) no-cue and in the dark (right) tracks. **h**, Running speed as a function of the position for nC1 (top), nC2 (middle), and dark (bottom) trials. Each heat map is normalized by its maximum speed.

### Distance coding in grid cells without visual cues

Grid cells can maintain periodic firing with a limited number of visual cues in 1D tracks^21^ or no cues^19,22,23^ without forming stable fields. However, it is unclear whether such unanchored distance coding can support an animal’s ability to navigate to the reward^23^. In line with these previous studies, we found that grid cells exhibited periodic activity in both the no-cue (Fig. 2e-f) and the dark tracks without forming stable fields along the track. The spatial period of this activity was ∼1.87 times larger than the grid scale measured in the two-dimensional real enclosure, scaled equally across all grid modules (Fig. 2g, Spearman correlation nC1: ρ=0.54, *P*=3.49 x 10^-55^; nC2: ρ=0.6, *P*=1.21 x 10^-76^; dark: ρ=0.52, *P*=8.22 x 10^-56^; ratio between VR and RE spatial period mean±s.d.: nC1: 1.86±0.74; nC2: 1.88±0.66; dark: 1.87±0.71). Importantly, we found that mice did not decrease their running speed in anticipation of the reward on the no-cue track (Fig. 2h), indicating that they were unable to accurately estimate their position without visual cues. This suggests that grid cell distance coding may not be sufficient to support accurate path-integration-based navigation in the absence of stable anchoring fields.

### A single visual cue promotes field formation with a cell-type-specific spatial distribution

Next, we introduced a single prominent 1m wide visual landmark starting at 2.5m to investigate how a spatial reference point closer to the reward would affect field formation and the mouse’s ability to estimate the goal location. The presence of the visual cue significantly increased the number of fields formed in all mEC cell types (Figs. 3a-b). The newly formed fields were mostly clustered at the start of the track and near the cue (Fig. 3c, Extended Data Fig. 7b), with different functional cell types preferentially encoding distinct parts of the cue. We used a cue score^4^ to quantify the overlap between the field and the cue (see Methods). The cue score of grid cells was significantly lower than all other cell types (Fig. 3d). Together with spatial cells, they showed a bias to form fields towards the end of the cue (Fig. 3c), as indicated by a significant positive shift in their cue score (Fig. 3e). Similarly to the no-cue track condition, grid cells encoded distance in the one-cue track (Fig. 3f-g, Spearman correlation for the one-cue track 1C: ρ=0.58, *P*=5.74 x 10^-69^, ratio between VR and RE spatial period mean±s.d.: 1.93±0.73).

**Fig. 3:**
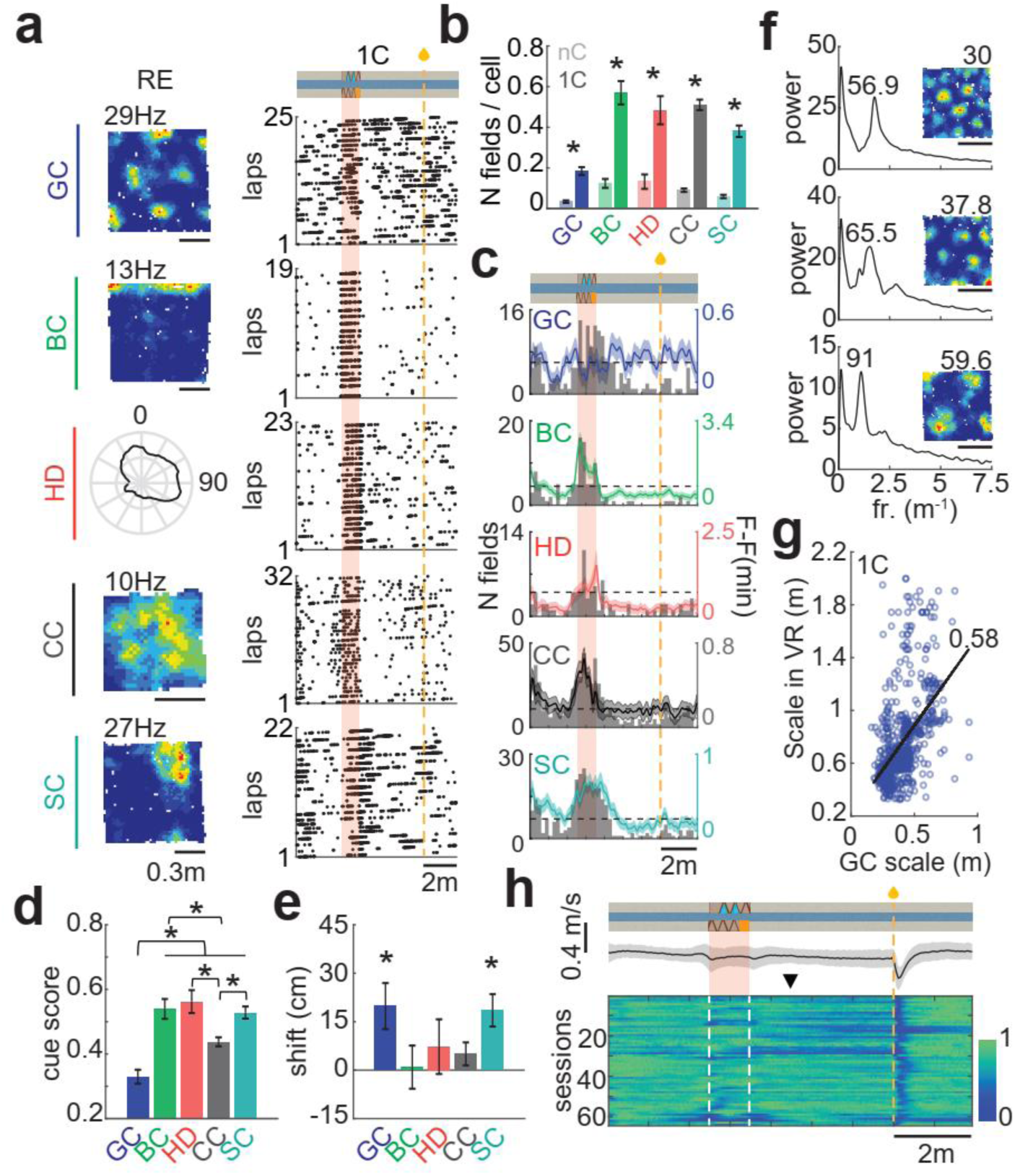
A single visual cue triggers the emergence of fields with a cell-type-specific spatial distribution. **a**, Typical rate maps and raster plots of different functional cell types in RE and in the one-cue (1C) VR track, respectively. **b**, The average number of fields (including cells with zero field) per cell type in the nC (lighter shade) was significantly lower than in the 1C (darker shade) track (Wilcoxon signed-rank test: GC z=-8.25, *P*=1.58 x 10^-16^; BC z=-6.88, *P*=6.01 x 10^-12^; HD z= -5.52, *P*= 3.32 x 10^-8^; CC z=-15.65, *P*=3.38 x 10^-55^; SC z= -11.05, *P*= 2.21 x 10^-28^). Grid cells were significantly less likely to make a field in the 1C track compared to all other cell types (Wilcoxon rank-sum test: BC vs. GC z=-9.68, *P*=3.84 x 10^-22^; HD vs. GC z=-6.02, *P*=1.73 x 10^-9^; CC vs. GC z=-11.29, *P*=1.49 x 10^-29^; SC vs. GC z=-6.93, *P*=4.15 x 10^-12^) **c**, Field count overlaid with the smooth firing rate in the 1C track (as in Fig. 2c). **d-e**, Average cue score (d) and corresponding spatial shift (e). The cue score was lower in grid cells compared to all other cell types (Kruskal-Wallis test with Bonferroni correction: HD vs. GC *P*= 2.8 x 10^-6^; SC vs. GC *P*=1.69 x 10^-9^; BC vs. GC *P*=7.57 x 10^-7^; CC vs, GC *P*= 0.0011). There was a significant shift in GC and SC (mean±s.e.m. spatial shift values, one-sample t-test: GC 19.82±7.13, t=2.78, df=109, *P*=6.4 x 10^-3^; BC 0.93±6.67, t=0.14, df=74, *P*=0.89; HD 7.28±8.46, t=0.86, df=45, *P*=0.39; SC 18.51±5.01, t= 3.69, df=184, *P*= 2.99 x 10^-4^; Wilcoxon signed-rank test: CC 5.04±3.55, z=1.22, *P*=0.22). Only cells forming at least one field were included. **f**, Typical Fourier spectrograms shown in Fig. 2e of different scale grid cells. The spatial period in VR (cm) in the 1C track (above the FFT power peak), increases with the grid scale. Grid scale (cm) indicated above the 2D firing rate maps. Scale bar: 50 cm. **g**, Grid scale in 1C VR track vs. its respective grid scale in 2D RE. **h**, Normalized average running speed in the 1C track. The black arrow indicates the position at which some animals begin to decrease their speed (∼1m after the cue), suggesting their anticipation of the reward from this point.

Only 2.2% (68/3160) of entorhinal principal cells could form a field between 4.5m and 6.5m. Notably, the running speed of the mice either did not decrease before the reward site or started to decrease from one meter after the cue at 4.5m location (Fig. 3h). The latter suggests that, similarly to the no-cue track, the mice may not have been able to accurately estimate their position starting from one meter after the cue onwards, where field formation collapsed. This range is consistent with the previously reported path-integration range^24^.

### Sublinear increase in the number of fields vs. cues in mEC cells

We further hypothesized that the abundance of visual cues in the cue-rich track would lead to enhanced field formation, given that the mice could accurately self-localize there (Fig. 1f). Indeed, the number of fields increased further in the cue-rich environment (Fig. 4a-c). However, it was consistently smaller than the number of available cues across all cell types, with grid cells showing the weakest ability to form fields in response to visual cues. This suggests that field formation may be constrained by a coordinated inhibitory cell network^8,25,26^ rather than following a simple stimulus-response rule.

**Fig. 4:**
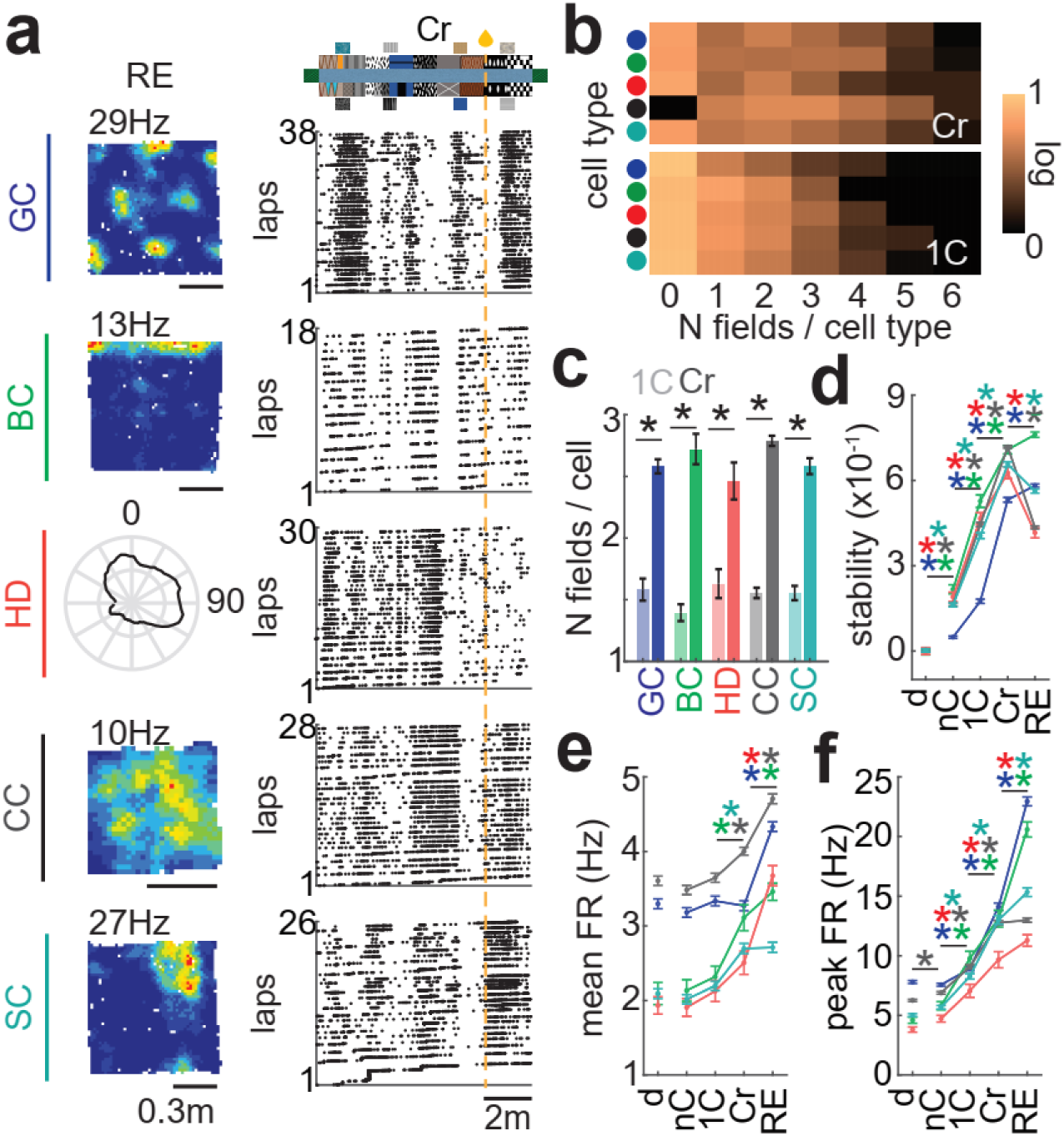
Sublinear increase of the number of fields per entorhinal cell as more visual cues are available. **a**, Typical rate maps (top left: peak firing rate) and raster plots of different functional cell types in RE and in the cue-rich (Cr) VR track, respectively. **b**, Distribution of the number of fields formed per cell for each cell type in cue-rich (top) and one-cue (bottom) track; color code as shown in (a). All pixel values have been converted to base-10 logarithms for display. **c**, Average number of fields (excluding cells with zero field) per cell type in the one-cue (1C, lighter shade) and cue-rich (Cr, darker shade) tracks (number of fields mean±s.e.m. 1C vs. Cr, Wilcoxon rank-sum test: GC 1.58±0.09 vs. 2.58±0.06, z=-8.18, *P*=2.89 x 10^-16^; BC 1.39±0.07 vs. 2.72±0.12, z=-7.17, *P*=7.43 x 10^-13^; HD 1.63±0.12 vs. 2.46±0.15 z=-3.74, *P*=1.87 x 10^-4^; CC 1.55±0.04 vs. 2.79±0.04, z=-16.14, *P*=1.27 x 10^-58^; SC 1.55±0.06 vs. 2.58±0.06, z= -9.45, *P*= 3.27 x 10^-21^). **d-f**, Average spatial stability (**d**), peak firing rate (**e**), and mean firing rate (**f**) measured in different VR tracks and RE for different functional cell types. The color code for the cell type is as in (a). The spatial stability increased in all cell types as more cues were added (stability mean± s.e.m. 1C vs. Cr, Kruskal-Wallis test with Bonferroni correction: GC 0.18 ± 7 x 10^-3^ vs. 0.53 ± 8.10^-3^, *P*=2.28 x 10^-119^; BC 0.53±0.022 vs. 0.71 ± 0.015, *P*=9.54 x 10^-6^; HD 0.47±0.02 vs. 0.63±0.021, *P*=3.44 x 10^-4^; CC 0.45 ± 0.007 vs. 0.72±0.004, *P*=5.53 x 10^-94^; SC 0.41±0.01 vs. 0.66±0.009, *P*=1.13 x 10^-47^). The peak firing rate increased in all cell types as more cues were added (peak firing rate mean± s.e.m. 1C vs. Cr, Kruskal-Wallis test with Bonferroni correction: GC 8.92 ± 0.16 vs. 14.12 ± 0.3, *P*= 8.76 x10^-37^; BC 9.78±0.59 vs. 13.11±0.71, *P*=0.0061; HD 7.06±0.54 vs. 9.67±0.68, *P*=0.011; CC 9.07±0.16 vs. 12.76±0.19, *P*=1.61 x 10^-54^; SC 8.27±0.23 vs. 13±0.36, *P*=4.81 x 10^-19^). The mean firing rate significantly increased in border cells, spatial cells and cue cells as more cues were added (average firing rate mean± s.e.m. 1C vs. Cr, Kruskal-Wallis test with Bonferroni correction: GC 3.34 ± 0.065 vs. 3.27 ± 0.067, *P*=1; BC 2.32±0.14 vs. 3.11±0.18, *P*=7.86 x 10^-3^; HD 2.13±0.14 vs. 2.51±0.16, *P*=0.96; CC 3.65±0.061 vs. 4±0.052, *P*=7.64 x 10^-7^; SC 2.2±0.062 vs. 2.7±0.068, *P*=5.54 x 10^-7^). Significant difference between consecutive datapoints is indicated with an asterisk using the same color code as for cell types.

The increase in field number was accompanied by higher spatial stability, as previously described in hippocampal cells^27^, and increased peak and mean firing rates (Fig. 4d-f). The highest mean firing rate was observed in the real environment across all cell types, reflecting the richness of available information there, including multiple sensory cues and vestibular inputs (Fig. 4f).

### Grid cells show the strongest reciprocal connections with the local interneural network

Next, we asked how interneurons might contribute to the network architecture governing functionally defined medial entorhinal excitatory cells. We defined interneurons based on the peak-to-trough latency of their waveforms (≤0.3 ms), as previously described^15,28^. To examine excitatory connections from principal cells onto interneurons (E-I), we used spike-time cross-correlograms from pairs of simultaneously recorded neurons (Fig. 5a). As previously suggested, a short-latency (≤5 ms) peak in the cross-correlogram indicates a putative monosynaptic excitatory connection^8,29–32^. We calculated the fraction of putative monosynaptic connections per functional cell type as mice navigated the cue-rich virtual track (Fig. 5b). In line with our hypothesis, we found that the relative number of projections from principal cells to interneurons was higher compared to relative projections from principal cells to other principal cells (Fig. 5c, Extended Data Fig. 8a, 410/48040 pairs (E-I) vs. 1188/288331 pairs (E-E), permutation test corrected for difference in the number of pairs: *P*=0), despite the overall lower number of interneurons compared to principal cells^33^. Grid cells showed the highest fraction of significant excitatory projections to inhibitory neurons (Fig. 5b, permutation test corrected for difference in pairs number: GC vs. BC *P*= 0.0015; GC vs. SC/CC *P*=0; GC vs. HD *P*=6 x 10^-4^, Extended Data Fig. 8b-d). Most grid cells with putative connections to interneurons had a low head direction score (mean±s.d. head direction score 0.077±0.047; Extended Data Fig. 2a), which was in line with previous findings showing that non-directional grid cells establish more connections to interneurons than conjunctive grid cells^8^. Importantly, in most cases, single interneurons received inputs from several different functional cell types (61/92, on interneurons receiving inputs from at least two neurons of different types, Fig. 5d) and sent inhibitory inputs to one or two cell types, mainly grid cells and cue cells (Fig. 5e-i). Taken together, this indicates that the mEC spatial map may indeed be shaped by a shared pool of inhibition, and grid cells may play a key role in building a cohesive map, as previously suggested using computational modeling^9,34–36^.

**Fig. 5:**
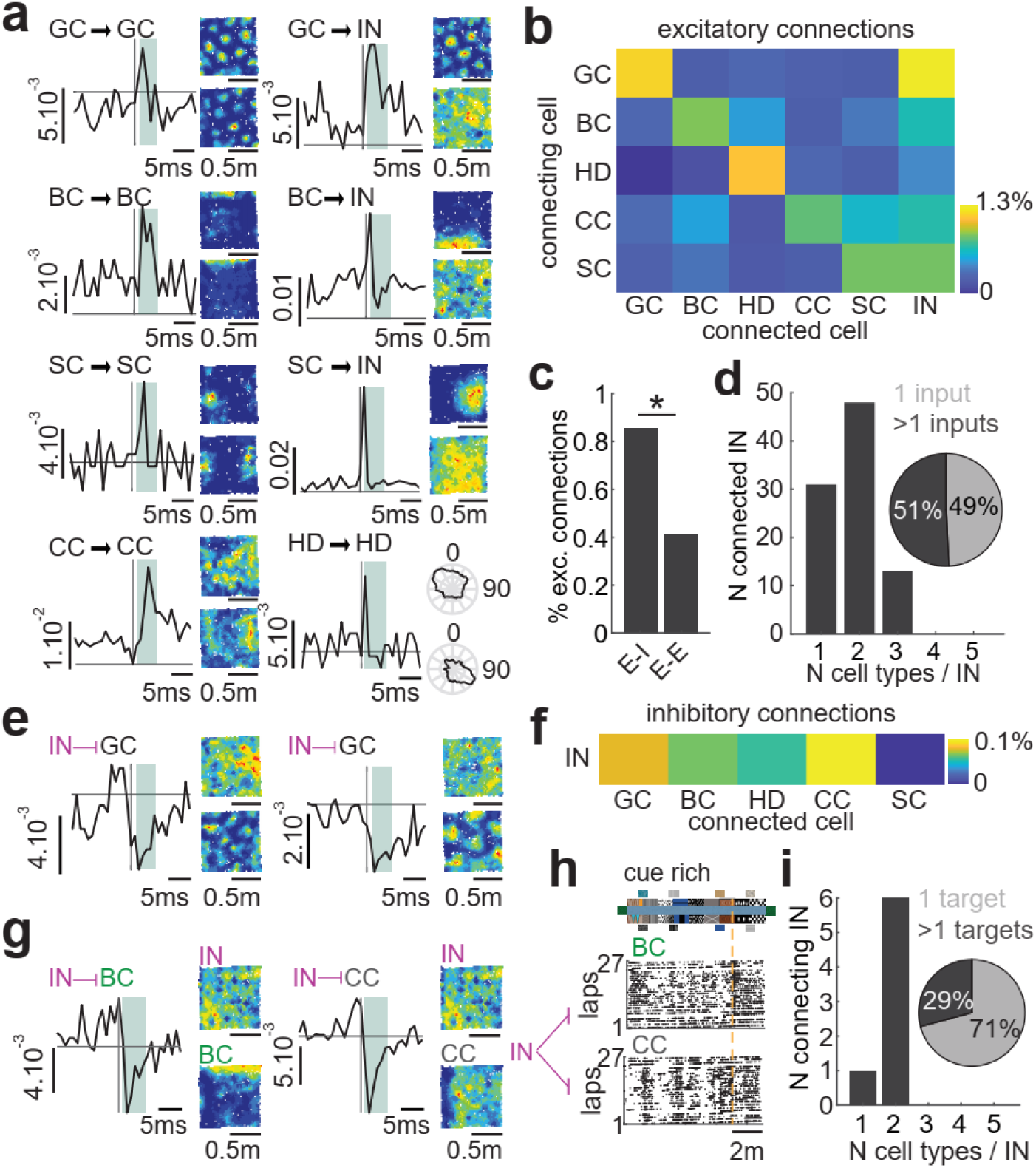
Grid cells show the strongest reciprocal connections with the local interneural network. **a**, Eight typical examples of cross-correlograms showing putative monosynaptic excitatory connections between two neurons with their rate maps shown on the right: a connecting (top) and connected (bottom) neuron. Cell types are indicated on the top left. The gray shaded area indicates the monosynaptic delay (1 to 5ms). **b**, Connectivity matrix of the percentage of monosynaptic excitatory connections (connecting neurons: y-axis; connected neurons: x-axis). Each bin indicates the proportion of significant connections among the total number of connections calculated for a given pair type. **c**, Percentage of monosynaptic excitatory connections for excitatory-to-inhibitory (E-I) and for excitatory-to-excitatory (E-E) neuron pairs. **d**, Pie chart: Percentage of interneurons connected (excitatory connection) by one (light gray) or more than one (dark gray) principal cell(s). Histogram: distribution of the number of different cell types connecting single interneurons (only interneurons connected by at least two principal cells are included). **e**, Two examples of inhibitory connections from interneurons to grid cells shown as in (a). **f**, Connectivity matrix as in (b) of the percentage of significant monosynaptic inhibitory connections per type of neuronal pair. **g**, An example interneuron (pink) sending inhibitory connections to both a cue cell (grey) and a border cell (green) in the cue rich track. Note that the two target cells were inhibited along the same left-side wall in the RE (bottom rate maps). **h**, Raster plots of the spiking activity in the cue-rich track of the CC and BC shown in (g). **i**, Pie chart: Percentage of interneurons connecting (inhibitory connection) one (light gray) or more than one (dark gray) principal cell(s). Histogram: distribution of the number of different cell types receiving inhibitory connections from single interneurons (only interneurons connecting at least two principal cells are included).

### Subnetwork connectivity by function and theta modulation

Given the crucial influence of theta oscillatory activity on the generation of grid cell patterns^37–39^, border cell acuity, and head-direction tuning^18^, we hypothesized that theta modulation may play a prominent role in field formation, and cells may show distinct connectivity motifs based on their function and theta modulation. To test this, we divided principal cells and interneurons into theta and non-theta categories, based on their theta scores^2^ (Fig. 6a-b). We found that in both populations theta-modulated cells tended to have shorter peak-to-trough latencies (Fig. 6c), and higher peak and mean firing rates (Fig. 6d-e). This suggests that theta- and non-theta cells may represent distinct neuronal subtypes.

**Fig. 6:**
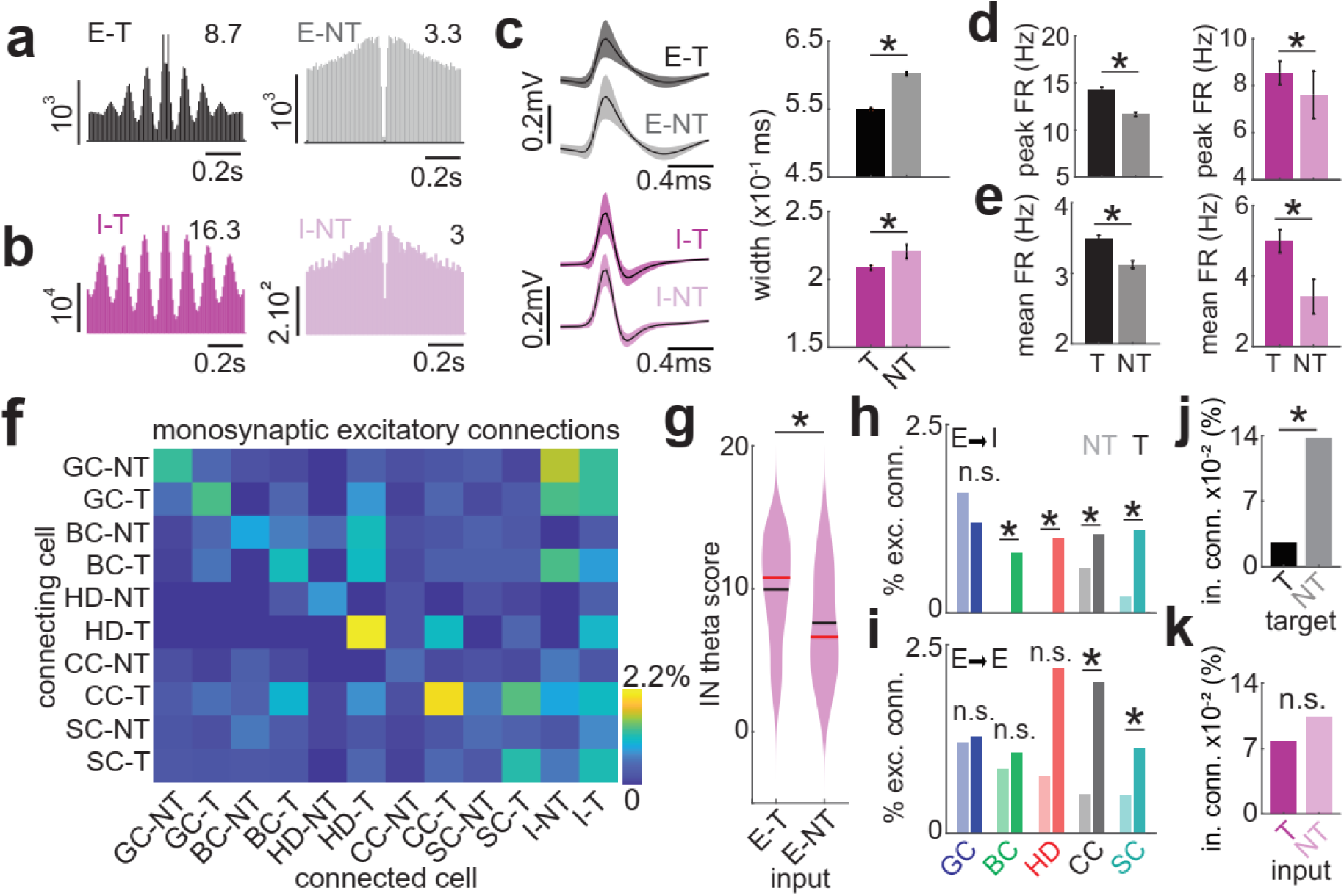
The recruitment of the inhibition is stronger in theta-modulated networks except for grid cells. **a**-**b**, Typical examples of auto-correlograms of T- and an NT-principal cells (E-T and E-NT) and interneurons (I-T and I-NT). Top-right: theta score. **c**, Mean ± s.d. of the spike waveforms (left) and mean ± s.e.m. of the waveform peak-to-through widths (right) for T- and NT-principal cells and interneurons (Wilcoxon rank-sum test waveform width interneurons T vs. NT: z=-2.21, *P*=0.0268; principal cells T vs. NT: z=-19.9, *P*= 3.61 x 10^-88^). **d-e**, Mean ± s.e.m. peak and mean firing rate for T- and NT-principal cells and interneurons (mean ± s.e.m. mean firing rate interneurons T vs. NT: 4.99±0.32 Hz vs. 3.42±0.49Hz, Wilcoxon rank-sum test: z=7.78, *P*=7.16 x 10^-15^; mean ± s.e.m. mean firing rate principal cells T vs. NT: 3.49±0.04Hz vs. 3.22±0.05Hz, Wilcoxon rank-sum test: z= 5.15, *P*=2.55 x 10^-7^; mean± s.e.m. peak firing rate interneurons T vs. NT: 8.54±0.49 Hz vs. 7.61±1.01Hz, Wilcoxon rank-sum test: z=5.62, *P*=1.9 x 10^-8^; mean ± s.e.m. peak firing rate principal cells T vs. NT: 14.33±0.22Hz vs. 11.97±0.21Hz, Wilcoxon rank-sum test: z=8.78, *P*=1.72 x 10^-18^). **f**, Percentages of putative monosynaptic excitatory connections as a function of the cell type and the theta category. **g**, Theta scores distribution (mean: red; median: black) of interneurons connected by either T- or NT-principal cells. **h**, Percentage of monosynaptic excitatory connections from T- or NT- (darker and lighter shade respectively) principal cells to inhibitory neurons of the same theta category. **i**, Same as (**h**) for excitatory-to-excitatory connections. **j**, Percentage of monosynaptic inhibitory connections from interneurons (of any theta category) to either theta principal cells (black) or non-theta principal cells (gray). **k**, Percentage of monosynaptic inhibitory connections from theta interneurons (T, dark pink) or non-theta interneurons (NT, light pink) to principal cells (of any theta category).

Next, we calculated the connectivity matrix of significant excitatory monosynaptic connections from principal cells to other principal cells and interneurons grouped by their functional type and theta modulation (Fig. 6f). The connectivity matrix revealed four major connectivity motifs (Extended Data Fig. 8e-f). In the dominant motif, as described above, the principal cells preferentially innervated interneurons over other principal cells. Moreover, the recruited interneurons tended to share similar theta affinity (Fig. 6g, Wilcoxon rank-sum test: z=5.89, *P*=3.89 x 10^-9^). In the second motif, across E-E connections between principal cells, principal neurons preferentially connected within their own cell type group and theta category, forming type-specific subnetworks, with consistent but weaker cross-subnetwork connectivity (Extended Data Fig. 8f). Border cells also showed additional strong connections to theta-modulated head-direction cells. In the third motif, projections between theta-modulated cell pairs dominated over non-theta pairs (Extended Data Fig. 8g). Grid cells were the only exception, showing comparable connectivity between theta and non-theta groups (Fig. 6h-i, permutation test corrected for difference in pairs number, excitatory-to-inhibitory: GC *P*=0.16; BC *P*=0.02; HD *P*=0.024; CC *P*=0.0094; SC *P*=3 x 10^-4^; excitatory-to-excitatory: GC *P*=0.57; BC *P*=0.63; HD *P*=0.28; CC *P*=0; SC *P*= 4 x 10^-4^). In the final ‘inhibitory’ motif (Extended Data Fig. 8h), non-theta principal cells tended to receive significantly more feedback inhibitory connections (Fig. 6j, permutation test corrected for difference in pairs number: *P*=0), despite theta-modulated principal cells tending to make more connections to interneurons compared to non-theta cells (motif 3). Theta and non-theta interneurons sent comparable amounts of inhibitory projections to excitatory cells (Fig. 6k, permutation test corrected for difference in pairs number: *P*=0.26). This suggests that the interactions between theta and non-theta streams may be implemented via an inhibitory network.

Notably, theta-modulated cue cells sent the strongest projections to all other cell types (permutation test, corrected for differences in pair numbers: *P*=0), potentially serving as a hub for transmitting sensory information about visual landmarks to other mEC cells. On the other hand, theta-modulated head direction cells were the strongest receiver of cross-type inputs, suggesting that they may act as a prime integrator and the mEC output hub. The strongest excitatory cross-subnetwork afferents to grid cells came from theta-modulated cue and border cells, which may provide information about external cues and the enclosure’s geometry, respectively, acting as reset and anchoring points to stabilize the grid, as previously suggested^40–43^.

## Discussion

We show that principal neurons in the medial entorhinal cortex operate via strongly interconnected subnetworks defined by their functional cell type and theta modulation. While these subnetworks have distinct neural architectures, they also share some striking common network motifs. The most dominant motif revealed greater relative connections from principal cells to a common pool of interneurons than to other principal cells. We hypothesize that this communication via common inhibition provides the basis of field formation in building a coherent spatial map, resulting in a sublinear field-to-cue relation. Furthermore, we showed that within the mEC map, theta-modulated cue cells are the prime hub for integrating external sensory information and conveying it to other cell types: their mean and peak firing rates, stability, and field-to-cue responses are most strongly associated with the number of cues, and they also provide the strongest projections to all other cell types. All cell types receive weak but consistent inputs from multiple other cross-type principal cells, thereby integrating exteroceptive and interoceptive signals and subsequently sending them to a shared inhibitory pool.

With the exception of grid cells, principal cells tend to connect to other principal cells and interneurons of similar theta category, with theta connectivity dominating over non-theta. This suggests that there may be two parallel streams encoding the same spatial variables. This is consistent with recent evidence showing that theta- and non-theta-modulated head direction cells respond differently to sensory changes^52^. Together, this points to a dominant theta-selective feedforward inhibitory loop, in which theta-modulated principal cells tend to recruit a shared pool of theta-modulated interneurons, and other theta-modulated cross-type principal cells. Theta oscillations may provide a time window for integration of sensory and locomotion information during navigation, while the non-theta stream may be involved in mnemonic information processing^14^. This hypothesis is in line with previous studies showing that layer II reelin cells, which send projections to the dentate gyrus and CA3, may be less theta-modulated compared to calbindin cells^44^ which project to CA1. The former projections were suggested to be important for signal retrieval, while the latter for active encoding during navigation^14^. Of all principal cells, grid cells show the strongest innervation with the inhibitory network, connecting the theta stream to the non-theta streams. Based on this, we speculate that grid cells may play a key role in coordinating the scene building into a cohesive map via inhibition, as previously suggested by the field-boundary-interaction model in rodents and bats^35,41,45^.

## Methods

### Subjects

Experimental procedures and animal use were performed in accordance with UK Home Office regulations of the UK Animals (Scientific Procedures) Act 1986, following ethical review by the University of Cambridge Animal Welfare and Ethical Review Body (AWERB). All animal procedures were authorized under Personal and Project licences held by the authors. Fifteen male C57BL/6J mice aged 2-6 months old were used as subjects. Mice were single-housed in cages on a reversed 12/12h light/dark schedule (with lights off at 9:00 AM).

### Surgical Procedure

Mice were anesthetized with 1–2% isoflurane in O2, and 0.05mg/10g body weight Metacam and chronically implanted with either tetrode micro-drives or Neuropixels 2.0 probes^17^. First, the skull was exposed, and four miniature screws were attached to the skull. One screw located to the left of the bregma served as a ground/reference electrode. A small craniotomy was performed above the posterior sinus, 3.4 mm from the midline. Next, the headplate was cemented. The mEC probes were implanted 3.2-3.4 mm lateral to the midline and 0.2-0.4 mm anterior to the transverse sinus, with an angle of 6 degrees in the anterior-to-posterior direction in the sagittal plane. Tetrodes were lowered in the brain at a depth of 0.6 mm. Neuropixels 2.0 probes were lowered in the brain at a depth of 2.5-2.85 mm. The microdrives were fixed to the skull with dental cement. After surgery, the mice were placed in a heated chamber for 1 to 2h until fully recovered from anaesthesia, then returned to their home cages. During the first 72 h after the surgery, the mice received Metacam (0.05mg/10g body weight) every 24 h. Mice were given 7 days of recovery with food and water ad libitum, after which the food restriction and behavioural training began. Four mice were implanted with eight tetrodes in the medial entorhinal cortex (mEC), eight were double implanted with eight tetrodes in left mEC and eight tetrodes in the right HP (the data not included in this study), and six mice were implanted with a Neuropixels 2.0 probe in the left mEC.

### Histology

At the end of the recordings, the mice were culled with an overdose of pentobarbital, transcardially perfused with phosphate-buffered saline (PBS), and then fixed in 4% paraformaldehyde. Brains were extracted and stored in 4% paraformaldehyde for at least 24 h. The brains were sliced at 30-50 microns in the sagittal plane. The brains implanted with tetrodes were stained with Nissl^46^ and the brains implanted with Neuropixels 2.0 were stained by immunochemistry targeting glial fibrillary acidic proteins (GFAP). For GFAP staining, the permeabilization solution used was PBS-Triton (PBS-T) x 0.25%, the blocking solution was 5% Normal Goat serum in PBS-Tx 0.1% (PBSGT), the primary antibody was a 1:500 dilution of rabbit anti-GFAP in PBSGT, and the secondary antibody was 1:500 Goat anti-rabbit 568 IgG (H+L).

### Data acquisition

mEC neurons were recorded as the head-fixed mice navigated in a one-dimensional virtual reality (VR) environment, followed by either one or two free-foraging 10-20-minute-long trials in a square-shaped arena (real environment, RE). The VR length was 9m, and the RE dimension was either 0.6 x 0.6 m or 1 x 1 m. The animals were directly transferred from the VR to the RE without disconnecting the pre-amplifier headstage, to record the same cells in both setups and classify the different cell types recorded in VR offline based on their spatial spiking patterns in RE.

### Virtual Reality Environment

We used our custom-built closed-loop 1D virtual reality (VR) environment, coupled with neural recordings from head-restrained mice as they ran linearly on a Styrofoam cylinder, facing two monitors displaying a visual scene controlled by the animal’s locomotion. The mice ran 9m-long laps and were teleported back to the start of the corridor at the end of each lap (the teleportation lasted 2 seconds). They received a milk reward at 7m on each lap. Two monitors were placed at a 90° angle to each other, in front of the mouse, to display the virtual scene generated by Unity Software. The rotation increments of the Styrofoam cylinder induced by the animal’s locomotion were read by a rotary encoder and sent via an Arduino Uno microcontroller (Arduino Encoder library version 1.4.0) to a custom-built LabVIEW program (v. 2018) that stored position and licking signal values at a 50 Hz sampling frequency. The milk rewards were delivered via a plastic tube positioned in front of the mouse’s snout. The delivery of the reward was controlled by voltage pulses sent from LabVIEW to a solenoid valve that functioned as a clamp on the plastic tube. During reward delivery, the tube was unclamped for 300 ms to dispense a drop of milk in front of the animal’s snout. Neural activity was recorded using a separate computer running either the OpenEphys system for tetrode recordings or the spikeGLX software for Neuropixels 2.0 probes. Synchronization TTL pulses were sent from the LabVIEW program to the electrophysiology recording system to align electrophysiological and position recordings offline.

### Real Environment

The real environment (RE) was designed to record neuronal activity from freely moving mice in a square-shaped arena (either 1×1m or 0.6×0.6m with a wall height of 0.5m). Two infrared light-emitting diodes (LEDs) were fixed on the animal’s head to track (x, y) position coordinates as well as the head orientation and direction of the animal, which were recorded and saved by a custom-built LabVIEW program. Synchronization TTL pulses were sent from the LabVIEW program to either the OpenEphys system for tetrode recordings or the spikeGLX software for Neuropixels 2.0 probes to align electrophysiological and position recordings offline.

### Behavioural training

The goal of the behavioural training was for mice to run on a virtual track to a reward while being head-fixed, and then to freely forage in the RE. After mice were trained on a cue-rich track, they were tested on the One Cue protocol (n=15 mice). During the first two days of habituation, the mice freely explored the cylinder, the head fixation bars, and the delivery reward tube of the VR setup. Over the next 2-4 days, the animals were trained to be head-fixed and to walk or run on the cylinder, with VR screens displaying a uniform dark background and a milk reward delivered frequently to encourage movement. The duration of head-fixation was progressively increased from 1-2 min to 20 min until the animal was comfortable on the wheel, ran the 9m track in < 1min, and consumed milk upon reward delivery. The mice were then trained to run on a 9m-long cue-rich VR track, with rewards delivered at 7m. The training lasted 4-6 days, during which the animals decreased their running speed at 1 to 1.5m before the reward location, showing that they could estimate their position relative to the reward location in the virtual track. After the cue-rich training was completed, the experimental protocol started. A typical session for the one-cue protocol consisted of 4 trials of 20-25 laps each. For each lap, the animal ran 9 meters, matching the length of the cue-rich track, and was then teleported back to the start. During teleportation, the screens displayed a uniform dark background for 2 seconds (inter-lap interval, ILI). Between VR trials, the screens were displaying a uniform dark for 2 min (inter-trial interval, ITI). The reward was delivered at position 7m for all laps except during the dark inter-trial interval. The four VR trials were: 1) a track with no visual cue available (no cue 1, nC1); 2) a track with one visual cue (1 m wide) on the track wall centred at 3m and spanning the whole height of the wall (1 cue, 1C); 3) a repeat of the track used in trial 1 (no cue 2, nC2), 4) the same cue-rich track used during training (cue-rich, Cr). The walls and floors of the no-cue and one-cue tracks had a random pattern. In addition, the virtual track had no end wall, so no looming visual signal would indicate that the end of the track was approaching as the animal progressed in it (see Extended Data Videos 1 and 2). After each VR recording session, the mice were recorded for 10-20 min in the RE to functionally identify the mEC spatial cell types that were recorded in VR.

### Spike extraction and spike clustering

Per session, the RE and VR recordings were concatenated prior to automatic spike sorting using Kilosort 2^17^. The output clusters were then manually curated using our custom-made Matlab program. Only spike clusters with a distinct waveform, a mean firing rate above 0.16 Hz and a refractory period ratio ≤0.8% were included to ensure high cluster quality. The refractory period ratio was calculated from the spike-time autocorrelation from 0 ms to 10 ms with a bin width of 0.5 ms. The refractory period ratio was defined as the sum of the spikes in the interval [0,1.5ms] in the auto-correlogram divided by the total number of spikes. Putative duplicate clusters were removed as described previously^4^. Briefly, the Pearson correlation coefficient between VR cue-rich trial smooth firing rate maps of cell pairs was calculated. If the correlation was above 0.8, the pair was classified as potentially overlapping. The cell with the lowest peak firing rate in the pair was removed from the analysis.

### Cell type classification

Spatial cell types were classified based on firing rate maps recorded in the real 2D square enclosure. The firing rate map was constructed by dividing the number of spikes fired in each 2cm × 2cm bin by the respective dwell time in seconds. The smoothed firing rate map was obtained by applying adaptive smoothing^46^. Five different cell types were classified based on their spatial properties: grid cells (GC), mEC spatially tuned cells (SC), which showed spatial tuning in the real 2D enclosure, head direction cells (HD), border cells (BC), and cue cells, which showed spatial tuning in the VR (Extended Data Fig. 1e).

Cells were classified as grid cells using the GC score^1^. Briefly, six local maxima were identified in the smoothed spatial autocorrelogram of the firing rate map as the six local maxima closest to the central peak (excluding the central peak itself). A mask on the spatial auto-correlogram, centred on the central peak but excluding the peak itself, was rotated in 30-degree increments up to 150 degrees. For each rotation, the Pearson product-moment correlation coefficient was calculated against the unrotated mask. The GC score was calculated as the difference between the minimum correlation coefficient for rotations of 60 and 120 degrees and the maximum correlation coefficient for rotations of 30, 90, and 150 degrees. A cell was classified as a grid cell if the GC score was ≥0.27^46^. The grid scale was defined as the radius of the circle, centred on the autocorrelation map that yielded the highest grid score.

Cells were classified as spatially tuned mEC cells if their spatial information (SI) score^47^ was ≥1.3. Cells were classified as head direction cells if their Rayleigh vector length was ≥0.19^46^. Cells were classified as border cells if the border score^3^ was ≥2. Briefly, cells were pre-selected as putative border cells if the proportion of bins positioned ≤10 cm from the wall with a firing rate ≥20% of the maximal firing rate was ≥50%. On this pre-selected set of cells, a mask with a value of 2 at each pixel on the surface of interest and 0 elsewhere was created. The border score was calculated as the sum of the normalized firing rate at the border divided by the sum of the mask value over the total number of elements in the mask. All grid cells, spatial cells and border cells that passed the score criterion were individually inspected and discarded if misclassified by the algorithm (false positives). Conversely, entorhinal cells not classified as a grid, spatial or border cell by the algorithm, but with clear spatial characteristics, were included (false negatives): about 20% of cells fell into these categories.

If cells were not classified in any of the cell types described above but had at least one spatial field formed in the VR cue-rich trial, these cells were classified as cue cells. Entorhinal cells were assigned to only one cell type, even when they scored high on more than one criterion, in the following order of priority: GC, BC, SC, HD, CC. For instance, if a cell met the criterion for both border and head direction cells, it was classified as a border cell and not a head direction cell.

We defined interneurons based on the peak-to-trough latency of their waveforms (≤0.3 ms), as previously described^15,28^.

### Firing rate, spatial stability and cue score

In VR, the unsmoothed rate maps were obtained by dividing the number of spikes per 5cm bin by the respective dwell time in seconds. The peak firing rate was calculated as the maximum value of the smooth rate map. The smooth rate map was obtained by applying the Gaussian kernel 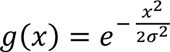 with σ = 1 and *x* ∈ {−1, 0, 1}. The mean firing rate was calculated as the total number of spikes per trial divided by the trial duration in seconds. In the real enclosure and in the VR, stability was defined using the Pearson correlation coefficient between the smooth rate maps of the first and second halves of the recording. The peak and mean firing rates were calculated as for the VR recordings.

The cue score was calculated as previously described^5^. Briefly, a ‘cue template’ was defined as a 1D vector of length 180 spatial bins (bin size = 5cm) with bin values equal to 1 where the visual cue is on the virtual track and 0 elsewhere. The spatial cross-correlogram between the cue template and the smooth rate map of individual neurons was calculated. The cue score was defined as the cross-correlogram peak (excluding local peaks with values below the average cross-correlogram value) corresponding to the spatial shift closest to 0.

The three 2-min-long dark inter-trial intervals were concatenated together for the dark track analysis. The annotation nC indicates an average between nC1 and nC2 data.

### Grid cells periodic firing in VR

To determine whether grid cells exhibit periodic firing in 1D VR in the absence of cues, we used Fourier analysis of individual cells’ inter-spike distance distribution. Only grid cells that fired more than 200 spikes during a VR trial were included in this analysis. First, the distribution of all pairwise inter-spike distances was calculated in centimetres. Then, the power spectrum of the distribution was calculated using the Fast Fourier Transform with padding to 2^13^ points for higher resolution (a sampling frequency equal to 100 points per meter). The power spectrum was smoothed by applying the Gaussian kernel 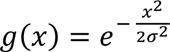 with *σ* = 5 and *x* ∈ {−10, −9, …, 9, 10} and peaks exceeding 10% of the maximum of the smoothed power spectrum were selected. Only power spectrum peaks corresponding to spatial periods between 30 cm and 200 cm were included in this analysis. When two peaks or more were detected, the one corresponding to the highest frequency was used, and when no peak was detected, the cell was discarded from the analysis.

### Defining spatial fields in VR

Spatial fields were defined as consistent, recurring spike firing in a defined region, with little activity outside that region. Only cells with mean firing rates between ≥0.16Hz and ≤8.3Hz and classified as spatial in VR (see below) were used for the field detection analysis.

Cells were classified as spatial in VR if their peak activity exceeded what would be expected by chance. To calculate the chance level, spike train and positions were offset 100 times by a random time interval ranging from 20 seconds to the trial duration minus 20 seconds, similar to what was previously described by Krupic and colleagues^46^. At each iteration, a surrogate smooth firing rate map was calculated and converted to a logarithmic scale. The chance level was defined as the 99^th^ percentile of the 100 maximal values (one per surrogate). A cell was classified as spatial if at least one spatial bin in the smooth rate map converted to logarithmic scale was superior to or equal to the chance level.

Next, to identify the individual fields, the spikes were first clustered using K-means (built-in MATLAB function ‘kmeans’) with the optimal number of clusters set to the number of peaks + 4 (a peak was defined as a local maximum above 30% of the peak in the smoothed firing rate). From this initial set, only clusters at positions corresponding to firing rates above the 60th percentile of the maximal values calculated from the surrogates and spanning at least 2 consecutive bins were defined as fields. If two adjacent peaks in the smooth rate map were closer than six bins (i.e., 30 cm), they were merged into a single field. Furthermore, the fields had to satisfy the stability criterion. Specifically, fields were defined as stable if the 60^th^ percentile of the values of lap pairwise correlation was ≥0.6 and if there were at least two spikes per lap on at least 50% of the laps in the field. To calculate the Pearson correlation between the laps, for each lap, the smooth rate map with only the spikes in the field was calculated.

### Theta modulation analysis

We calculated the theta modulation score of individual neurons recorded on the VR cue-rich track using the Fast Fourier Transform power spectrum of the spike-train autocorrelograms (calculated on a 500-ms window)^2^. The theta score is defined as the mean spectral power within 1 Hz of each side of the peak in the 6-10 Hz frequency band, divided by the mean spectral power between 2-125 Hz. A cell is defined as theta-modulated if the theta modulation score is≥5.

### Connectivity matrix

To determine the putative monosynaptic connections between mEC cells, we computed cross-correlograms for all the co-recorded pairs of the identified cell types (GC, SC, HD, BC, CC, and IN). A cross-correlation peak value above the chance level within a synaptic time window indicates that one cell is significantly firing near the time the other cell is firing, and thus these cells may be connected^28^. Temporal cross-correlations were computed using a custom Python program and the PyTorch library for GPU-accelerated computation. For each cell of the pair, its spiking activity was stored in a vector with a bin size of 1ms containing a value of 1 if at least one spike was emitted in that bin or zero otherwise. Cross-correlograms were generated on 30-ms time windows by calculating the Pearson correlation coefficient between two spike vectors (from two co-recorded cells) for shifts incrementing in 1-ms increments. If the maximum value in the resulting cross-correlogram was detected in one of the synaptic windows defined as [-10ms, 0ms] or [0ms, 10ms], then two test steps were performed to determine whether the peak is significant or not. The first step consisted of randomizing the indices of the spiking vector of one of the cells in the pair and running the correlation step described above against the non-randomized spiking vector of the other cell in the pair. The shuffling procedure was repeated 50 times per cell pair. The shuffle threshold was defined as the median value of the fifty 99^th^ percentile values of the shuffled cross-correlograms. If the peak in the pair cross-correlogram was above the shuffle threshold in either the [-10ms, 0ms] or the [0ms, 10ms] window, then the second significance test was performed: half of the indices of one neuron spike train were randomly chosen and cross-correlated with the whole spike train of the other neuron of the pair to obtain a half cross-correlogram. Then, the same half of the indices in the first neuron were randomized and cross-correlated with the whole spike train of the other neuron to get the shuffled half cross-correlogram. The peak of the half cross-correlogram was compared to the median of the values in the shuffled half cross-correlogram. This operation was repeated 10 times, and if the peak in the half cross-correlogram was above the shuffle threshold at least 9/10 times, then the pair was classified as significantly connected. Monosynaptic connections were defined as the pairs with cross-correlogram peaks occurring within 5-10 ms time windows. Negative correlations were detected following the same procedure, using 1^st^ percentile instead of 99^th^ percentile in the first significance test and multiplying the cross-correlogram values by (-1) before comparing to the median value in the second significance test.

### Statistical analysis

Matlab R2019a and Python 3.12 were used to perform basic statistical analyses. The standard error of the mean (s.e.m.) was used unless stated otherwise.

To compute the chance level of the percentage of cells forming a field in the no-cue track as a function of the number N of cells per cell type group, N numbers were drawn from the uniform distribution on the interval [0,1] over 500 iterations, and the 95^th^ percentile was calculated.

To compute the chance level of the field position distribution, field positions were randomly allocated to position bins (bin size=5cm), and the maximal number of fields per bin was stored. This was repeated for 500 iterations, and the 95^th^ percentile of the stored values was calculated.

For repeated-measures data that are not normally distributed (e.g., the number of fields per cell as the animal is exposed to different virtual tracks with varying numbers of cues), the Kruskal-Wallis test was used. When the dataset was normally distributed (e.g., running speed across tracks), one-way ANOVA was used. When pairs of datasets were compared using either the Kruskal-Wallis test or one-way ANOVA, the Bonferroni correction was applied. The normality of the dataset was checked using the MATLAB built-in function ‘kstest’.

To compare the same set of cells between two different tracks (e.g., stability, mean/peak firing rate, or number of fields per cell, including zero field), the Wilcoxon signed-rank test was used (for non-normally distributed and paired data).

To compare sets of cells across two tracks (e.g., the number of fields per cell, excluding zero fields), the Wilcoxon rank-sum test was used (for non-normally distributed data).

To compare the frequency of occurrence of an event between two groups of the same size (e.g. the percentage of cells of one cell type that would form fields or have synaptic connections to other cells in two different tracks), a permutation test was used. Specifically, the two datasets were shuffled using the MATLAB built-in function ‘randperm’, and the difference in the number of occurrences between the two shuffled datasets was calculated. This was repeated for M=5000 iterations, and the final p-value was calculated as the number of times the absolute value of the difference between the two unshuffled datasets was less than the absolute value of the difference between the shuffled datasets, divided by M.

To compare the occurrence frequency of an event between two groups of different sizes, N and (N-k), the permutation test was used with two down-sampling steps. First, to compute the difference of occurrences between the unshuffled groups, the larger group was randomly down-sampled to size (N-k) and the absolute difference was calculated, then averaged over 100 iterations. Second, to compute the difference of occurrences between the shuffled groups, the larger dataset was similarly down-sampled to size (N-k) before computing the difference at each of the M shuffling iterations. The differences were effectively computed between groups of size (N-k). Specifically, the two datasets were shuffled using the MATLAB built-in function ‘randperm’ and the difference in the number of occurrences between the two shuffled datasets was calculated. This was repeated for M iterations, and the final p-value was calculated as the number of times the absolute value of the difference between the two unshuffled datasets was less than the absolute value of the difference between the shuffled datasets, divided by M. To compare the proportion of cells with a field in the no-cue track across cell types, M=5000; for all tests performed in the connectivity analysis, M=10000.

## Data availability

Electrophysiological recording data generated in this study will be made available on the Zenodo data depository upon publication. Source data will be provided with this paper.

## Code availability

The codes for replicating the analyses in this study will be deposited on GitHub upon publication.

## Acknowledgments

We thank John O’Keefe for comments on the manuscript, Tereza Vargova for help with pilot experiments, Charlie Clarke-Williams for sharing the initial version of the cross-correlation script, Bogdan Toader for optimization of the cross-correlation script and Marino Krstulovic for help with manual curation of the cells.

This work was supported by the UK Dementia Research Institute, through UK DRI Ltd, principally funded by the Medical Research Council, by funding from the Cure Alzheimer’s Fund (J.K.). J.K. is a Wellcome Trust/Royal Society Sir Henry Dale Fellow (206682/Z/17/Z). The funders had no role in study design, data collection and analysis, decision to publish, or preparation of the manuscript.

## Authors information

### Contributions

Conceptualization: J.K. and P.K., Methodology: P.K., M.B. and J.K., Experiments: P.K. with contributions from J.K, Data analysis: P.K. with contributions from J.K. and M.B., Funding acquisition: J.K. Writing—original draft: J.K. Writing—review and editing: J.K., P.K., M.B.

### Corresponding authors

Correspondence to Pauline Kerekes or Julija Krupic.

---

**Extended Data Fig. 1:**
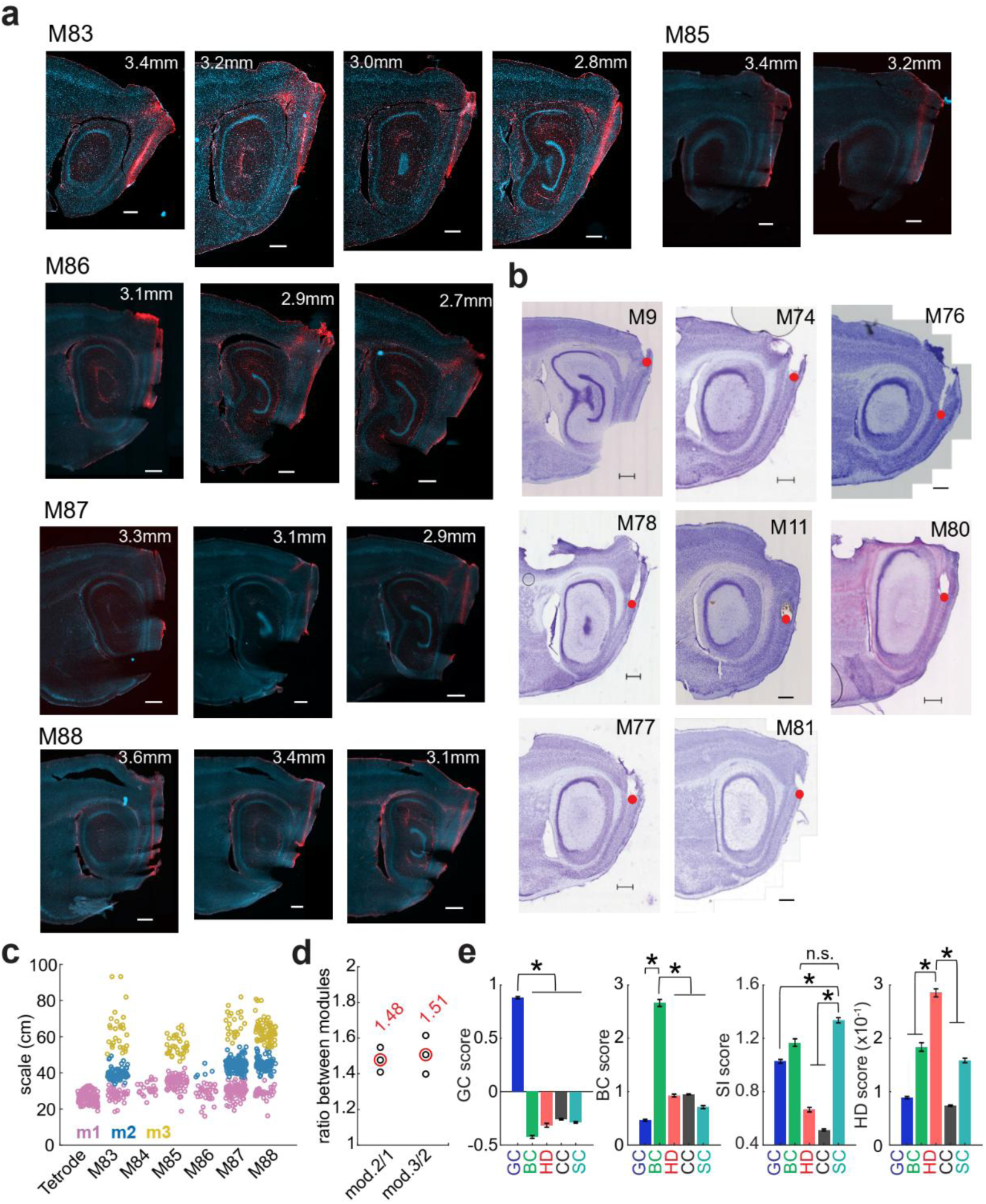
Recording probes, anatomical location, and cell type classification scores. **a-b**, Sagittal sections of the left mEC stained for GFAP (Alexa Fluor 568, red) and cell nuclei (DAPI, blue) in five mice implanted with Neuropixels 2.0 (a, each section shows a different Neuropixels shank; left-most is putative shank 0) and Cresyl violet for tetrode implants (b). The number above each section indicates the estimated medio-lateral position. The scale bar for (a) and (b) is 0.5 mm. **c**, Distribution of the grid scales across mice and clustering (using k-means) in three modules (m1 to m3). Mice implanted with tetrodes are pooled together (left) and Neuropixels animals (M83-88) plotted separately. **d**, Ratio between the average GC scales of two adjacent modules for the animals M83, M87 and M88 (black dots for single animals; red dots for the average). **e**, Average grid (GC), border (BC), spatial information (SI) and head direction (HD) scores for the five cell types. GC had the highest grid score (GC vs. BC/HD/CC/SC P<0.001), BC the highest BC score (BC vs. GC/HD/CC/SC P<0.001), BC and SC had a similar SI score value (BC vs. SC P=0.33) and both had the highest SI score (BC vs. GC/HD/CC P<0.05; SC vs. GC/HD/CC P<0.001), HD had the highest HD score (HD vs. GC/BC/CC/SC P<0.001, Kruskal-Wallis test with Bonferroni correction).

**Extended Data Fig. 2:**
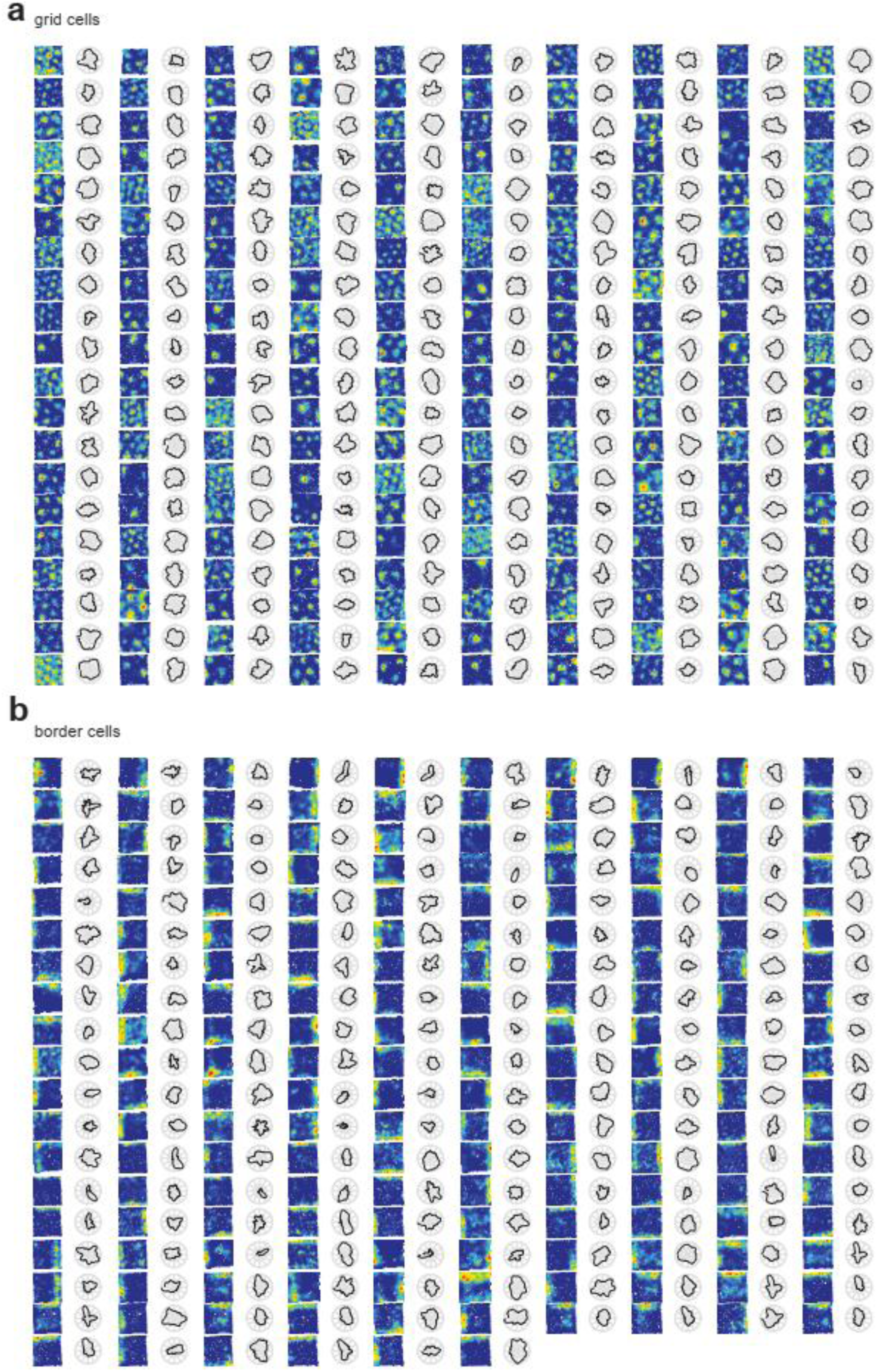
Example grid and border cells activity in the real enclosure. **a-b** Example of randomly selected grid cells (a, N=200/944 cells) and border cells (b, N=186/186 cells) spatial activity in the real enclosure (RE) as rate maps (left) and head direction tuning as polar plots (right).

**Extended Data Fig. 3:**
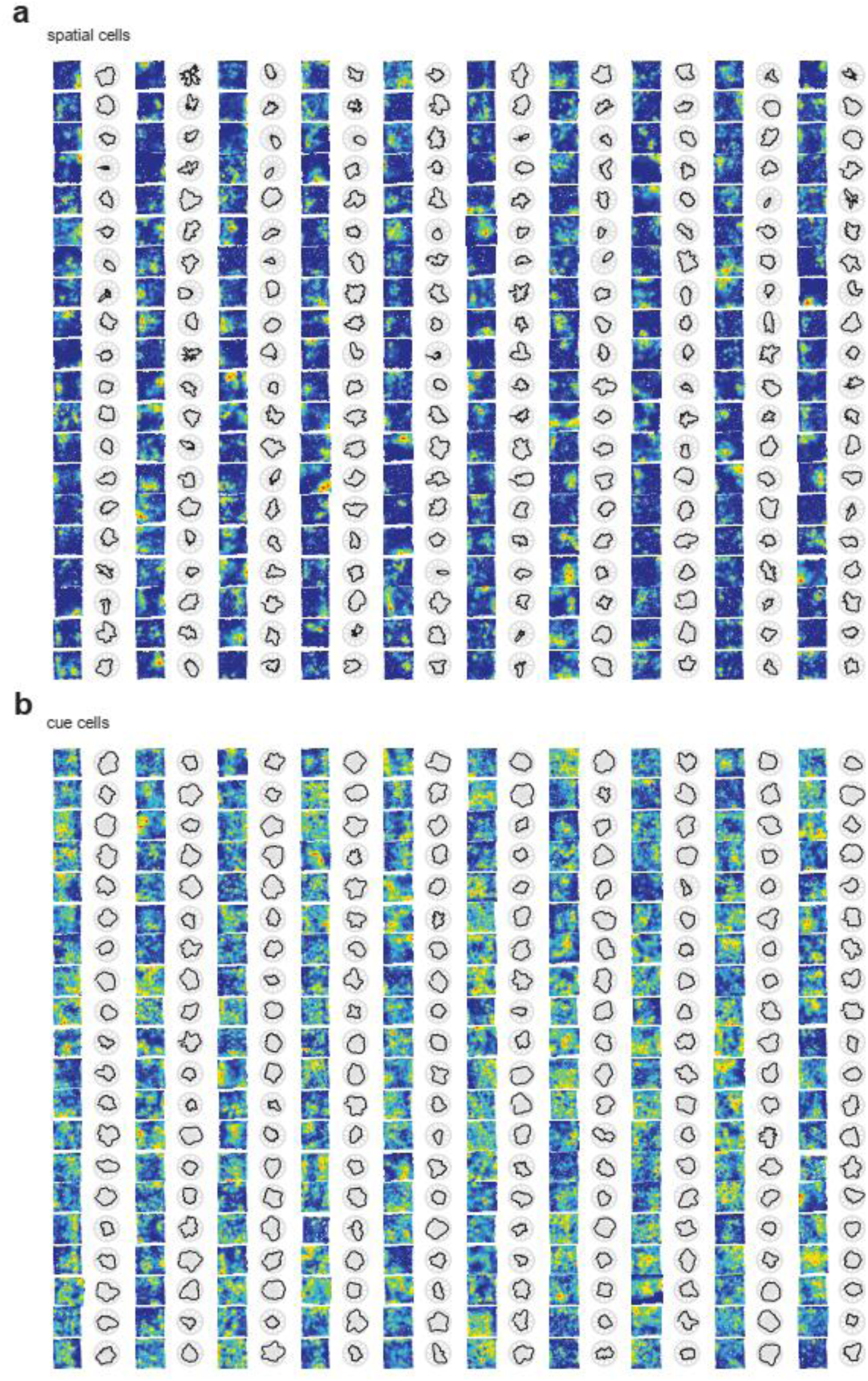
Example spatial cells and cue cells activity in the real enclosure. **a-b** Example spatial cells (a, N=200/760 cells) and cue cells (b, N=200/1115 cells) spatial activity in the real enclosure (RE) as rate maps (left) and head direction tuning as polar plots (right).

**Extended Data Fig. 4:**
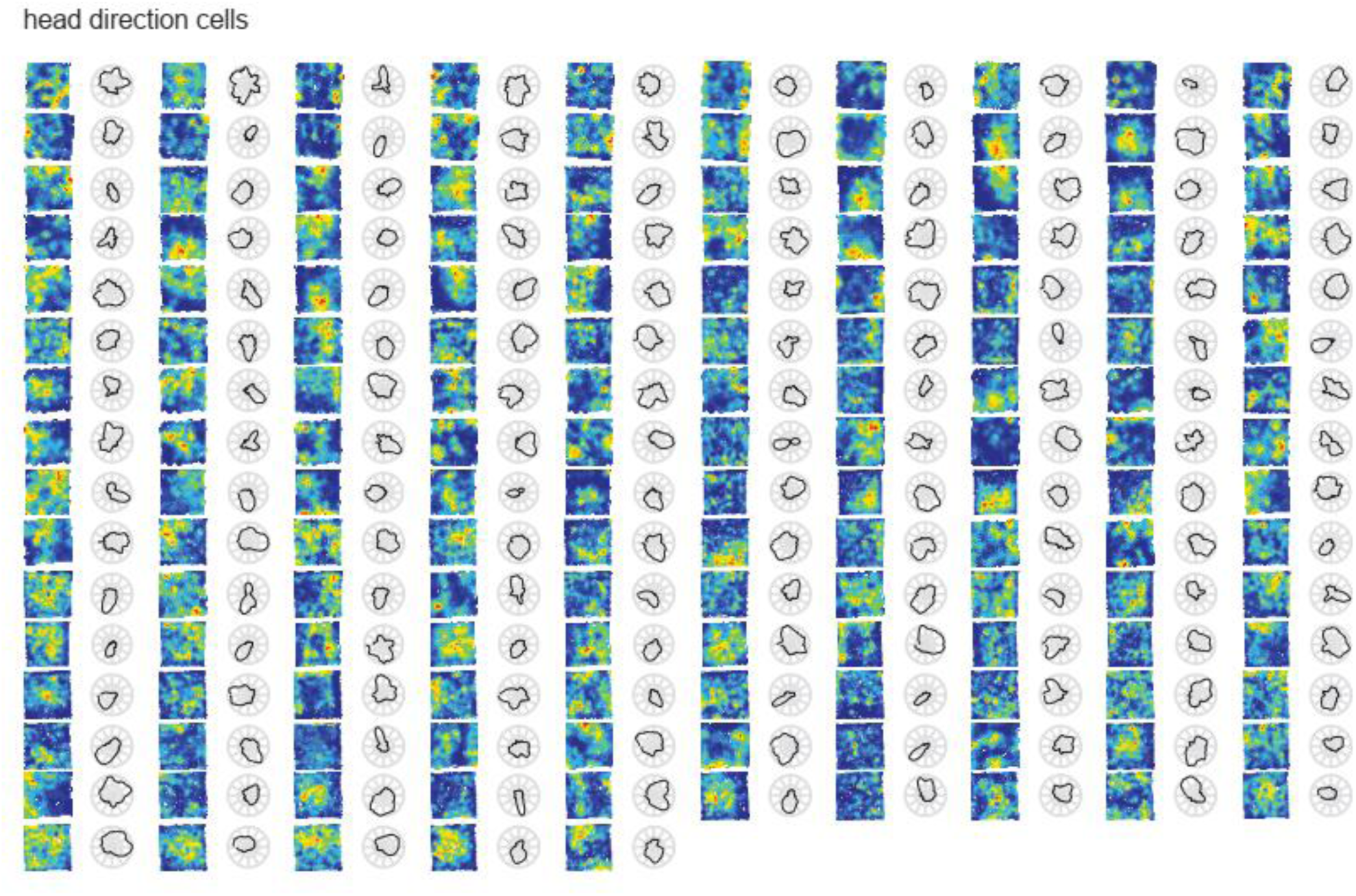
Example head direction cells activity in the real enclosure. Example head direction cells (N=155/155 cells) spatial activity in the real enclosure (RE) as rate maps (left) and head direction tuning as polar plots (right).

**Extended Data Fig. 5:**
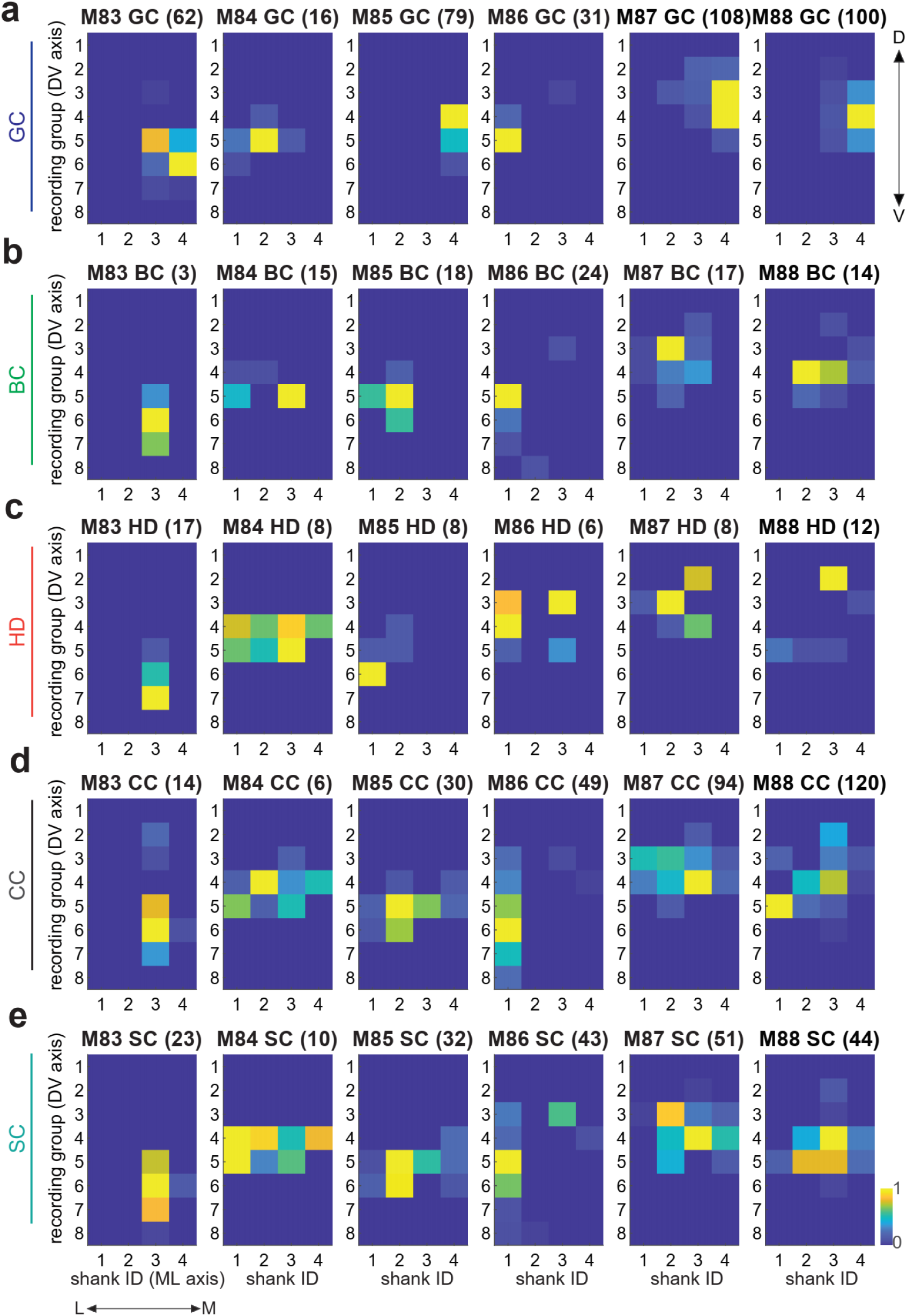
Distribution of the cell types on the Neuropixels 2.0 shanks. **a**, Anatomical distribution of grid cells (GC) on the four shanks of the Neuropixels 2.0 probe for individual mice (M83 -88). Each bin corresponds to a recording group. The y-axis and x-axis show the dorsal-to-ventral and lateral-to-medial axes, respectively. The number above each heatmap is the maximum number of cells per recording group. Each heat map is normalized by its maximal number of cells / recording group. **b-e**, same as (a) for border, head direction, cue, and spatially tuned cells.

**Extended Data Fig. 6:**
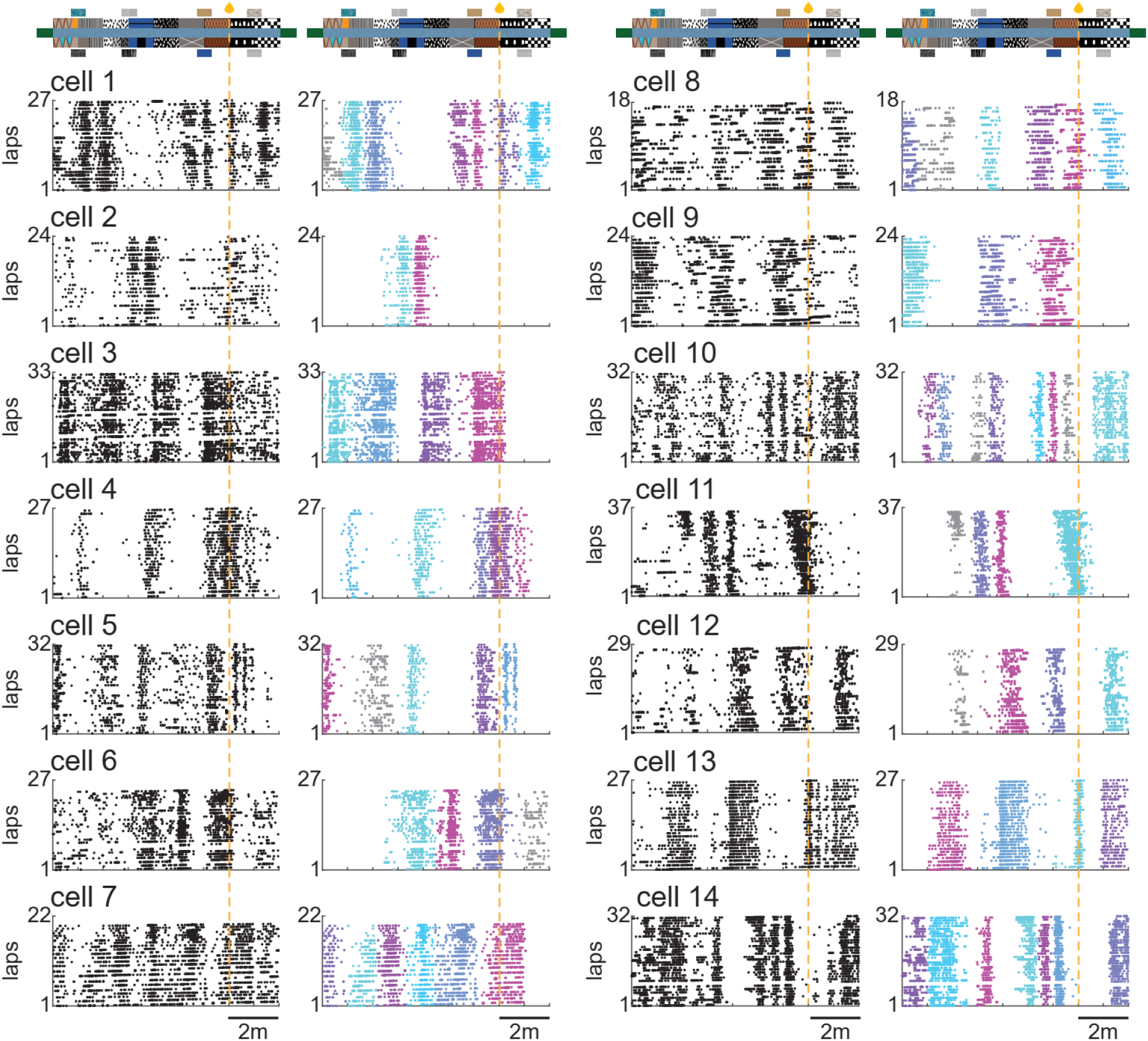
Examples of the field detection algorithm output. Fourteen typical examples of raster plots of different neurons recorded in cue-rich VR tracks (left, black dots showing spikes) with identified fields shown in different colors (right). The unstable fields are shown in gray and have been excluded from the analysis. A field was classified as unstable if in-field spikes were present on less than 50% of the laps (example cell 11) and/or if the lap-to-lap stability was too low (example cells 6 and 8). Eyeballing inspection confirmed a correct field identification >80% of the time.

**Extended Data Fig. 7:**
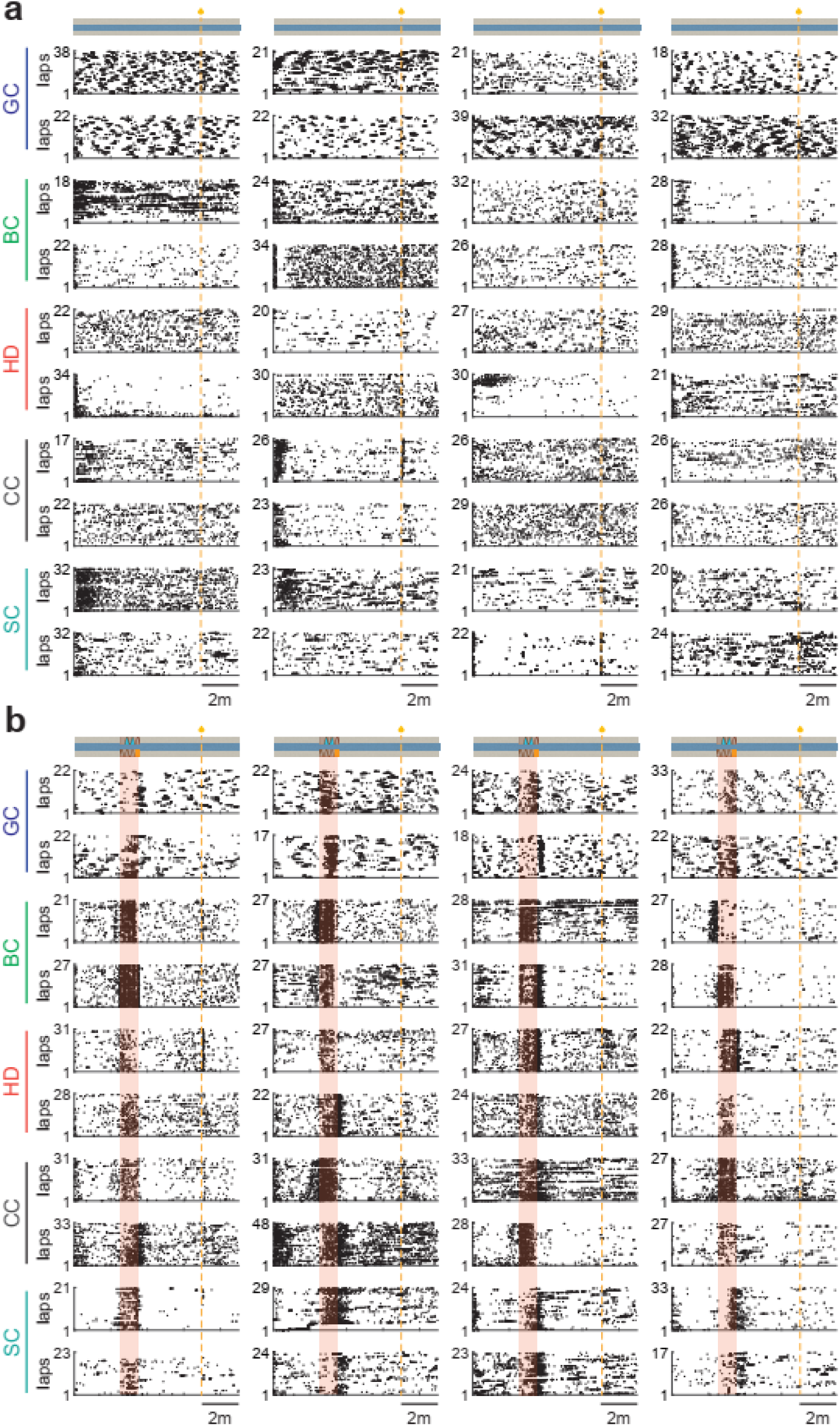
Example cell activity in the no-cue and one-cue virtual tracks. **a**-**b**, Forty-eight typical examples of grid (GC), border (BC), head direction (HD), cue (CC) and spatially tuned (SC) activity in the first trial of the no-cue (a) and in the one-cue (b) VR tracks. The cell type is indicated on the left (eight different cells per cell type). The golden vertical dashed line indicates the location of the reward and the shaded area the cue location in (b).

**Extended Data Fig. 8:**
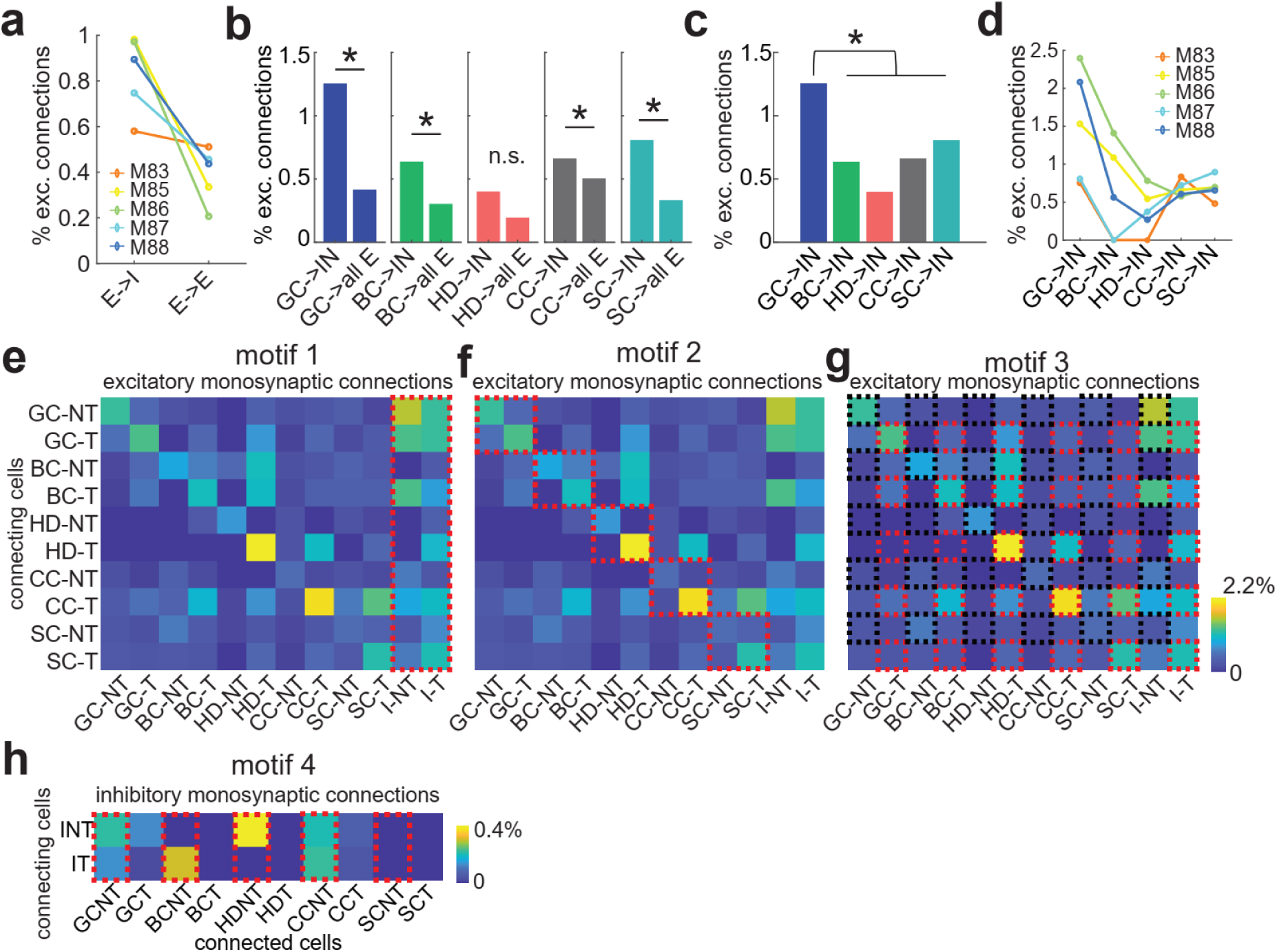
Characterization of the excitatory and inhibitory connections as a function of cell types and theta categories. **a**, Percentage of putative monosynaptic excitatory-to-inhibitory connections for individual mice. **b**, The percentage of monosynaptic excitatory-to-inhibitory is higher than excitatory-to-excitatory connections for all cell types except HD (permutation test corrected for unequal number of pairs: *P*=0 for GC and SC; *P*=0.0119 for BC; *P*=0.0023 for CC; *P*=0.0973 for HD). **c**, Comparison of the percentage of monosynaptic excitatory-to-inhibitory per cell type (GC sends more inputs to inhibitory neurons than all other cell types). **d**, Percentage of monosynaptic excitatory-to-inhibitory connections per cell type for individual mice. **e-h**, connectivity motifs in mEC neurons: Motif 1: a higher number of projections from principal cells to interneurons (red rectangle) than to other principal cells (e); Motif 2: a higher number of projections between cells that belong to the same cell type (f); Motif 3: higher number of projections between theta-modulated (red) than between non-theta-modulated (black) cells (g); Motif 4: non-theta modulated cells receive more inhibitory projections than theta modulated cells (h).

